# Metabolic blueprints of monocultures enable prediction and design of synthetic microbial consortia

**DOI:** 10.64898/2026.01.11.698878

**Authors:** Sarah Bald, Rowan Nelson, Jing Zhang, Markus de Raad, Dillon C. Scott, Kendall Hughes, Melisa Osborne, Ilija Dukovski, Trent R. Northen, Daniel Segrè

## Abstract

Synthetic microbial ecology aims at understanding complex microbial dynamics and designing communities with desired properties, often based on mathematical models of individual organisms. However, it is unclear whether simplified models harbor enough detail to predict the composition of synthetic communities, especially in metabolically complex environments. Here, we use longitudinal exometabolite data of monocultures for 15 rhizosphere bacteria to parametrize a consumer-resource model, which we use to predict pairwise co-cultures and higher order communities. The capacity to artificially “switch off” cross-feeding interactions in the model demonstrates their importance in ecosystem structure. Leave-one-out and leave-two-out experiments reveal that pairwise co-cultures do not necessarily capture inter-species interactions within larger communities and broadly highlight the nonlinearity of interactions. Finally, we illustrate that our model can be used to identify new sub-communities of three strains with high likelihood of coexistence. Our results highlight experimental and theoretical challenges to be addressed in future efforts and establish hybrid mechanistic and data-driven metabolic models as a promising framework for predicting and engineering microbial communities.

## Introduction

Synthetic microbial communities are a powerful and rapidly expanding avenue for studying aspects of natural microbial ecosystems, such as dynamic environments and microbial interactions, while maintaining tractability of laboratory settings ^1–6^. An open question is whether the study of synthetic communities could lead to the discovery of underlying rules of community assembly and help rationally steer communities towards desired states, with applications in agriculture ^7,8^, environmental sustainability ^1,9^, waste management^10,11^, metabolic engineering ^12,13^ and biomedicine ^14,15^. Efforts to predict community assembly and dynamics in either synthetic or natural microbial communities require appropriate mathematical models capable of capturing the metabolic properties of individual microbes and their interactions.

Microbial interactions are often defined and measured based on co-culture experiments, in which the effect of one organism on another is directly assessed. While powerful, this approach is generally difficult to extend to higher-order interactions and to the prediction of whole communities, as additional species can rewire competition and cross-feeding networks, introducing nonlinear and context-dependent effects^16–18^. Alternatively, one can hypothesize that microbe-microbe interactions emerge from the metabolic activity of individual species dynamically altering the nutrient availability for others through consumption or production ^19,20^. Thus, ecosystem dynamics could be predicted based on the metabolic properties of individual organisms, i.e. microbe-environment (rather than microbe-microbe) interactions. These metabolic exchanges for individual species can be assessed by measuring extracellular metabolites over time, providing estimates of organism-specific uptake/secretion rates. These rates can then be used to infer the dynamics of communities of varying composition and complexity.

Plant-associated microbiomes are a compelling system to study in this context, since their central interactions are metabolite-mediated, their functions carry clear agricultural stakes, and standardized synthetic communities now make them tractable in the laboratory ^8^. These are complex ecosystems whose emergent functions remain poorly understood but hold strong potential for sustainability, including biofertilization and plant resilience under high temperatures and droughts ^21,22^. Plants enrich specific microbial communities by releasing carbon products from photosynthesis through their roots, while rhizosphere communities can supply usable nitrogen and induce pathogen resistance^23,24^. One notable standardized switchgrass-derived community can be reproducibly assembled from isolates, maintains stable diversity, and colonizes model grasses under laboratory conditions ^2,25,26^. This community can also be cultivated in a defined medium (Northen Lab Defined medium (NLDM)) that contains many root exudates and soil metabolites ^27^, enabling controlled, host-free studies of interspecies interactions and resource-use dynamics in a chemically identifiable environment.

The availability of a rhizosphere synthetic bacterial consortium that grows reproducibly on a defined medium offers the opportunity to develop efficient, scalable models for mechanistic predictions of ecosystem dynamics ^20,28–30^. Mathematical approaches for modeling microbial ecosystems span a range of complexity, from pairwise interaction frameworks like generalized Lotka-Volterra models ^29,31,32^, to detailed metabolic reconstructions like genome-scale metabolic models (GEMs) ^33^. Simpler models permit analytical derivations ^34–36^ and can capture complex behaviors, but they poorly account for nutrient feedback, environmental fluctuations, and context-dependent interactions ^16,34^. Conversely, GEMs explicitly represent metabolite-mediated interactions, but are challenging to construct for non-model organisms and to scale to multi-species communities. Their community-level use has therefore concentrated on human-associated systems, where high-quality models exist for many taxa ^37,38^, and remains limited for environmental isolates. Occupying a middle ground, microbial consumer-resource models (CRMs) have emerged as a flexible, semi-mechanistic framework ^39,40^. Although CRMs lack stoichiometric detail, they describe an effective metabolism for individual consumers, making it possible to test the microbe-environment interaction paradigm described above. These interactions are represented as uptake and secretion matrices, allowing competition and cross-feeding to emerge as time-dependent properties of each organism’s metabolic activity in a shared pool of resources.

A central challenge in implementing CRMs lies in parameterizing them to reflect metabolic traits of different strains and specific environmental chemical compositions ^41^. Most CRM implementations utilize statistical distributions of parameters rather than direct measurements, to capture general features of competition and cross-feeding. These statistically parametrized CRMs of arbitrary species and environments have been shown to successfully predict community-level phenomena, including large-scale ecological patterns observed in microbiome data ^39^, convergence on shared carbon sources ^40,42^, and complex community responses to environmental stimuli ^20,43–45^. However, their lack of specificity limits their use in rational community design, where predictive control requires parameterization linked to specific metabolites and organism-level metabolic traits ^28,41^. Prior work has used metabolomics-parametrized CRMs to explain features of specific communities, but detailed species-specific uptake and secretion matrices have not been encoded from a diverse, chemically defined medium.

Here we parametrize a CRM using longitudinal exometabolomics from monocultures of 15 switchgrass-derived isolates from the model synthetic community introduced above ^2,25^ to predict the dynamics and assembly of synthetic communities on NLDM (Fig. 1) ^27^. The parametrized CRM recapitulates monoculture behaviors and predicts outcomes and putatively exchanged metabolites in experiments across a range of complexity from co-cultures to entire community assemblies (Fig. 1d). The CRM reproduces the observed patterns of strain coexistence and community composition, while leave-one-out and pairwise removal perturbations uncover nuanced, nonlinear and context-dependent inter-species interactions (Fig. 1e). Finally, CRM predictions enable rational design of sub-communities with enriched diversity, highlighting that mechanistic, experimentally informed models can guide both understanding and engineering of microbial community dynamics (Fig. 1e).

**Figure 1.**
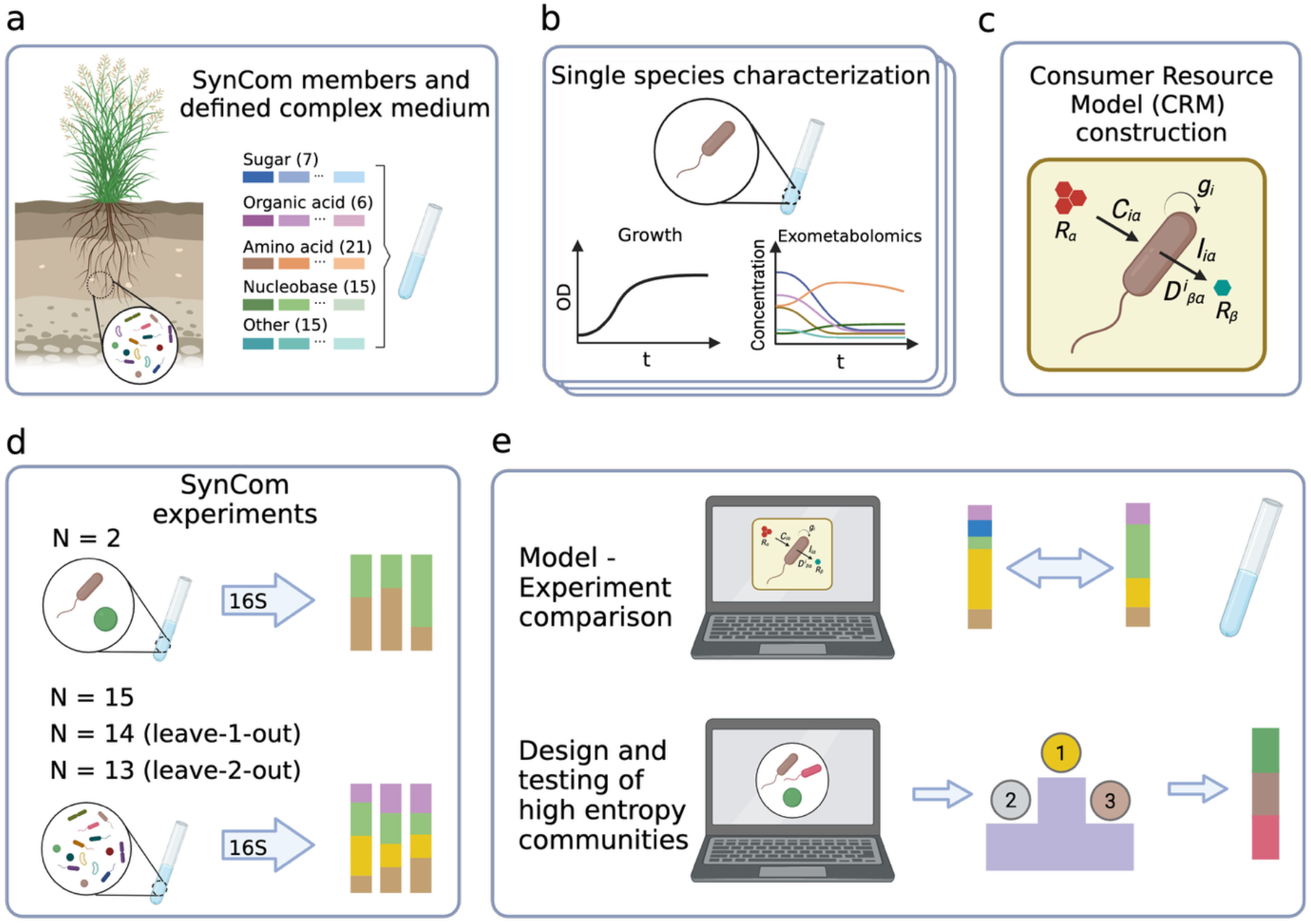
Workflow for data collection, model parametrization and testing of rhizosphere synthetic communities. **a**) The 15 microorganisms used in this study were isolated from the soil surrounding a switchgrass plant, *Brachypodium distachyon* (Coker et al. 2022). The isolates or communities were grown in a defined medium (NLDM) for cultivation and exometabolite profiling of soil bacteria (Supplementary Table 2). **b**) Isolates were grown in batch for 3 days, where growth yields were read using a spectrophotometer every 20 min, and metabolites in the spent media were sampled intermittently when cells were roughly grown at 25%, 50%, 75% and 100% of the final growth yields (Supplementary Fig. 1,2). **c**) Schematic of consumer resource modeling and model prediction of sub-community assembly. Organisms consume resources to produce biomass, meanwhile leaking metabolic byproducts into the shared environment, which can then be utilized by other co-cultured organisms for growth (Methods). Key parameters in the consumer resource models including C_iα_, D^i^_βα_, g_i_ were derived from the experimental data collected from monoculture. **d**) For model performance validation, communities containing 2, 15, or 14 (leave-1-out) strains were grown and passaged, and the final composition was determined by 16S sequencing. **e**) Consumer resource models fitted from monoculture were used to predict synthetic community abundances and design sub-communities with high entropy.

## Results

### Rhizosphere bacteria display complex and diverse metabolic behavior

We parameterized our ecosystem model by collecting monoculture growth and exometabolite data for each of the 15 rhizosphere bacteria (Supplementary Table 1). Each strain was cultivated on Northen Lab Defined Medium (NLDM) (Methods, Supplementary Table 2) to measure its growth curve and its consumption and secretion of 64 metabolites (amino acids, nucleobases, sugars, organic acids, vitamins). Metabolite utilization was quantified by collecting spent media for exometabolomic profiling across 4 timepoints, when cell densities roughly reached 25%, 50%, 75% and 100% of the final biomass, to quantify the net secretion/uptake of these metabolites within each interval (Supplementary Fig. 1-2). Strains differed significantly in their growth curves, with some strains *(Rhodococcus*, *Rhizobium*, *Arthrobacter)* growing to high yield after a short lag phase, and others barely displaying growth relative to negative controls (Supplementary Fig. 2). Notably, even the slower growers, *Methylobacterium* and *Marmoricola*, displayed measurable changes in medium composition relative to negative controls, suggesting substantial metabolic activity.

Clustering strains by their utilization of metabolites (Fig. 2a) reveal clear patterns across both strains and metabolites. *Variovorax*, *Methylobacterium*, *Bosea* and *Paenibacillus* are major secretors (over 18% metabolites with net increase) with major secreted compounds for these strains being sugars (Supplementary Fig. 3b). Moreso, *Variovorax*, *Methylobacterium* and *Bosea* each secrete more carbon than consumed at certain time points (Supplementary Fig. 3a). In contrast, *Burkholderia*, *Rhizobium*, and *Mucilaginibacter* depleted over 60% of available resources to levels 20-fold below their original concentration by the final harvest. We additionally observe that overall growth yield correlates well with total carbon uptake (r=-0.769, p=8e-4; Supplementary Fig. 3d). Overall, amino acids were the most broadly utilized metabolites across all bacterial strains, consistent with their established roles in plant exudates as signaling molecules and growth-promoting substrates that shape rhizosphere colonization ^46,47^. Despite these patterns, many resource utilization strategies are strain-specific (Supplementary Fig. 4). For 86% of metabolites (55/64), the range between the most and least active strain exceeded 1 (the size of the normalized range from complete depletion to no depletion of a metabolite), highlighting the breadth of metabolic diversity in the rhizosphere, and setting the stage for dynamic interactions during community assembly.

**Figure 2.**
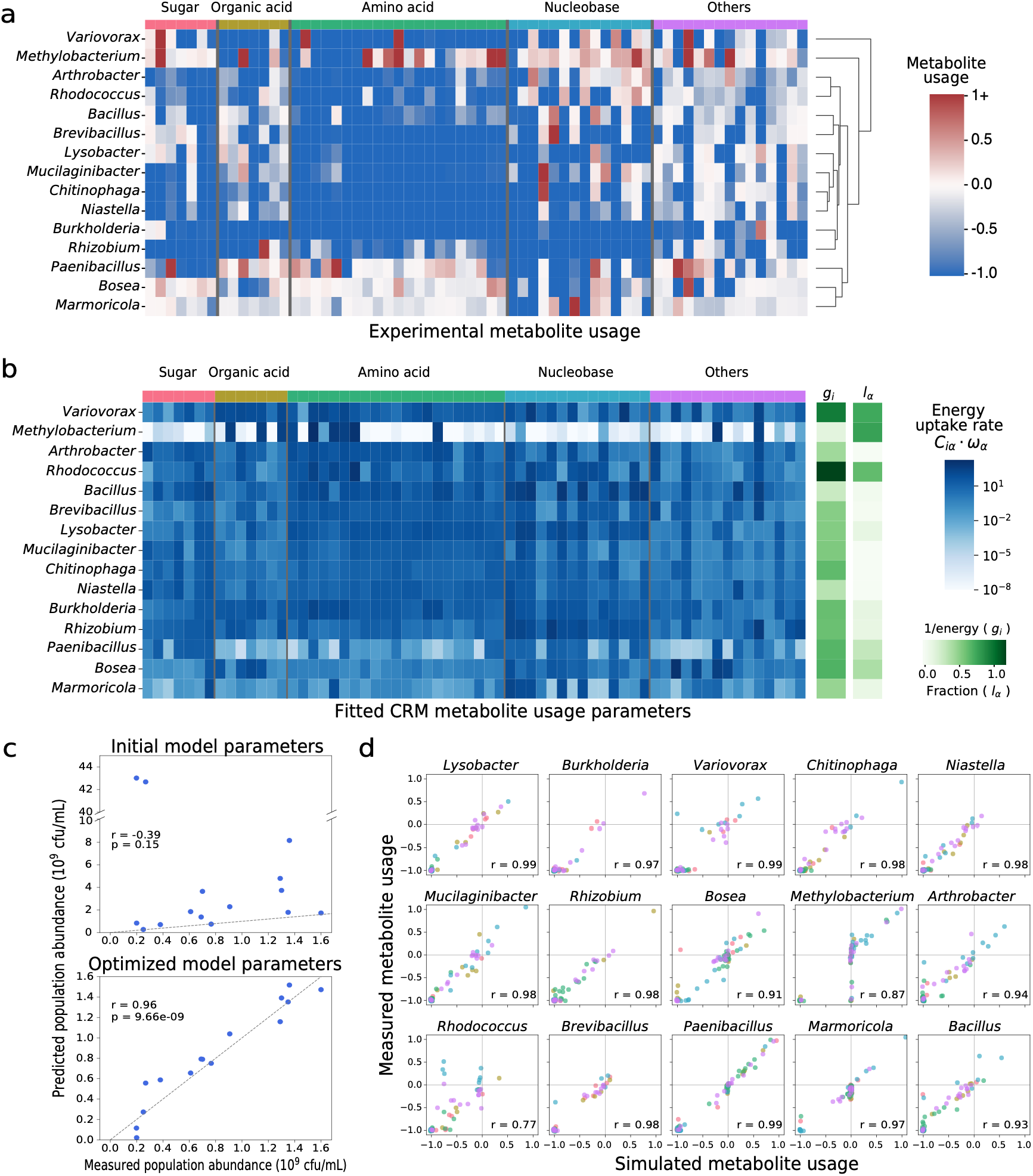
Parameter inference for consumer resource models using exometabolomic profiles and growth measurements from monoculture. **a)** Heatmap demonstrating the resource utilization and secretion by individual organisms in monoculture when growth reached maximum yield. Values (by colors) represent the abundance of each metabolite detected, with respect to its abundance in the blank media at that sampling point. A value above 0 (shown in red) means a net secretion of the particular metabolite and a value below 0 (shown in blue) means a net consumption. **b)** Parameters in the consumer resource models inferred from monoculture experimental measurements and further optimized by simulated annealing. Colors representing values of g_i_, l_iα_, and C_iα_*w_α_ are scaled according to the color bars. C_iα_*w_α_ values represent the metabolite uptake rate of a single CFU per unit of metabolite concentration (g/mL), expressed as energy acquired per cell per hour. **c)** Concordance of population abundance of each monoculture at the final sampling point determined by calibrated growth yield measurements, and predicted by consumer resource modeling with initial (top, n=15, Pearson’s r =-0.39) and optimized parameters (bottom, n=15, Pearson’s r = 0.96). A perfect fit 1:1 line is shown in grey. **d)** Comparison of resource utilization and secretion at the final sampling point determined by exometabolomics profiling and by model prediction for individual strains. A positive value denotes resource production, and a negative value denotes resource consumption normalized by metabolite initial concentrations. Different classes of molecules, including sugars, organic acids, amino acids, nucleobases and others are shown in corresponding colors as in Fig. 2a and 2b including sugars, organic acids, amino acids, nucleobases and others are shown in corresponding colors as in Fig. 2a and 2b. Pearson correlation coefficients are shown and each correlation has a sample size of 64 NLDM measured metabolites.

### Fitting the consumer-resource model from monoculture data

Growth curves and longitudinal exometabolite data from monocultures of each of the 15 strains were used to construct an experimentally parametrized CRM. Although the parametrization was performed at the level of individual strains, the resulting optimized parameters allow simulations at the ecosystem level, based on the assumption that interspecies interactions are mediated through metabolite consumption and production. The key inferred parameters are the resource consumption matrix (C) and the matrix describing how consumption of resources relates to exudation of others (D). Since C and D together encode consumption as well as exudation, cross-feeding is implicitly represented in the model, emerging from the exometabolomics data itself rather than from any additional dataset or interaction term. We fit these matrices from experimental data for each strain rather than sampling these parameters from predefined distributions ^39,40,42^ (Fig. 1b,1c). These experimentally derived metabolic preference parameters for each strain allow us to explicitly test whether monoculture behavior predicts community structure and dynamics in multispecies communities.

Our parameter inference procedure involves two steps. First, we derived estimates of strain-specific C and D matrix elements by fitting monoculture growth and time-resolved usage profiles of 64 metabolites to analytical model expressions (Methods, Supplementary Fig. 5). Unlike other CRM implementations, we treat D matrices as organism-specific, reflecting differences in intracellular metabolic capabilities in addition to resource preferences. By comparing how much each metabolite’s concentration changed between timepoints and assuming growth is proportional to consumption, the model equations simplify enough to solve directly for each strain’s consumption (C matrix), exudation (D matrix), and growth parameters (g_i_) (Supplementary Methods 1.2). These initial estimates, however, yielded model predictions that often predicted greater metabolite depletion relative to experiments (Pearson’s r =-0.39, Fig. 2c, Supplementary Fig. 6a). To address this, we performed a second optimization using simulated annealing benchmarked against individual growth and resource-use data (Methods). Parameters were iteratively perturbed to minimize error between simulations and experimental monoculture data (Supplementary Methods 1.3), revealing that while initial estimates generally overestimated growth, metabolite uptake and transformation rates (Supplementary Fig. 6), only moderate changes were made, with the exception of slow-growing strain *Methylobacterium* (Supplementary Fig. 6c-d). With optimized parameters (Fig. 2b), agreement between simulated and experimental growth yields substantially improved (Pearson’s r = 0.96, Fig. 2c), and metabolite utilization was accurately captured for most species (Pearson’s r > 0.9) (Fig. 2d). Only *Rhodococcus* (r = 0.77) and *Methylobacterium* (r = 0.87), exhibited comparatively weaker correlations. Importantly, both the analytical estimation and subsequent data-driven optimization proved to be jointly essential for quality parameterization of the CRM.

### Widespread coexistence in co-cultures recapitulated by parametrized CRM

To examine how the 15 strains interact in NLDM, we assembled all 105 pairwise co-culture communities. Strains were inoculated at equal OD abundance and passaged every 48 hours for 6 days, and final composition was measured by 16S amplicon sequencing (Fig. 1d). Additionally, these data were used to estimate pairwise interactions, defined as the overall effect of each strain on each other (Methods, Supplementary Fig. 7a).

Overall, coexistence (defined as the dominant strain composing <90% of final composition) was the most frequent outcome (68/105 pairs) (Fig. 3a). This likely reflects the high carbon concentration and metabolite diversity of NLDM, which supports multiple niches associated with metabolic preferences. Only two strains were consistently outcompeted by other strains: *Methylobacterium* grows much slower than other strains, and *Marmoricola* utilizes only a small set of NLDM metabolites. Organisms that often dominated without excluding partners (*Chitinophaga*, *Rhizobium*, *Bacillus*) are not necessarily the ones that grow fastest. We hypothesize that these strains are the ones that can benefit the most from interactions with others, possibly mediated by metabolite secretions. Conversely, fast growers generally performed well but were not consistently dominant, and intrinsic growth rate was a poor predictor of abundance in co-culture (r = 0.147, Supplementary Fig. 8a).

**Figure 3:**
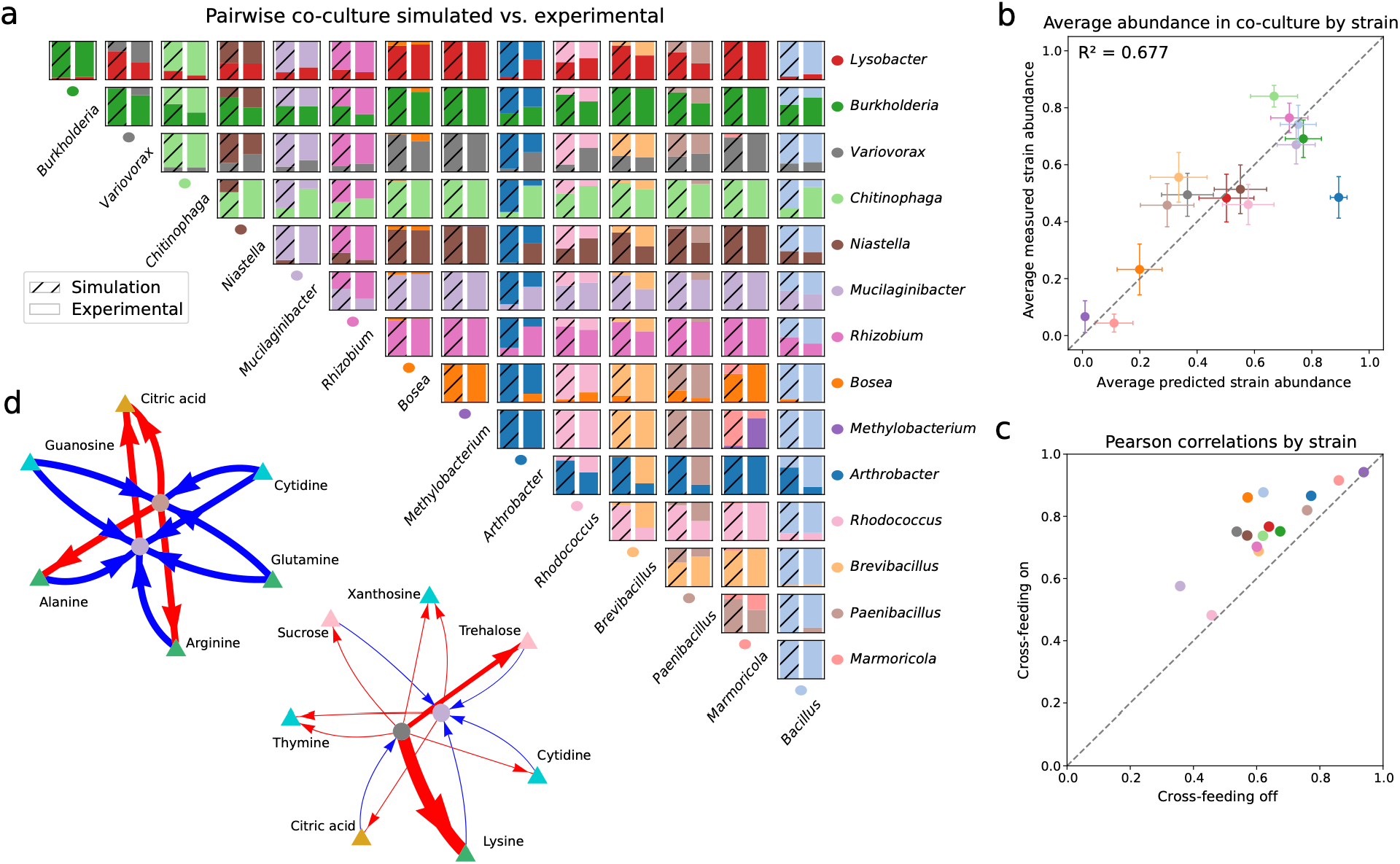
Pairwise co-culture modeling recapitulates community dynamics and elucidates key strain-metabolite interactions. **a)** Co-culture experimental and simulated outcomes using experimentally-informed consumer resource models. Each strain is represented by a unique color. The un-patterned bars show relative abundance of strains in experimental co-cultures, and the patterned bars show the predicted co-culture abundances from the CRM. **b)** Average strain abundance across 14 co-culture experiments predicted by the model compared to average measured strain abundance from 16s data, summarizing information from (a). Standard error bars are shown. Strain colors are consistent with (a). This same plot with each co-culture pair plotted individually can be found in Supplementary Fig. 8b. **c)** The predictive power (Pearson’s coefficient, n=14) of using consumer resource models with or without cross-feeding enabled in recapitulating community composition in co-cultures. Strain colors are consistent with A. **d)** Interaction networks for *Mucilaginibacter*, with *Variovorax* and *Paenibacillus*, showing significant cross-feeding of amino acids that benefits *Mucilaginibacter*. Production arrows are red and consumption arrows are blue, with widths relating to the rate of consumption or production determined from exometabolomic profiling. Comprehensive interaction networks of other strains can be explored interactively in the supplementary materials. Strain node colors are consistent with (a), and metabolite node colors are consistent with metabolite classes in Fig. 2a.

This dataset allowed us to test if a CRM parametrized solely from monocultures could explain pairwise co-culture assembly. By solving the CRM for all pairs (Methods), we generated an *in silico* dataset that was directly comparable to experiments (Fig. 3a, patterned bar plots). Predicted strain abundances in co-cultures display limited but statistically significant correlation with the corresponding measurements (R^2^ = 0.416, p=0.0002, Supplementary Fig. 8b). Interestingly, deviations from experiments are dominated by a few pairs involving *Arthrobacter*, which is often overpredicted. However, the CRM manages to accurately capture the average abundance of each strain across all its co-culture pairs (R^2^ = 0.743, Fig. 3b).

Beyond assessing accuracy, the CRM enables quantitative probing of the role of metabolite sharing in community dynamics. In the CRM it is possible to artificially “switch off” the metabolite exudation, disabling cross-feeding between strains (Methods). Switching off metabolite exchange decreased the predictive power of the CRM for all co-cultures (except pairs with *Methylobacterium* and *Marmoricola,* that were minimally affected), overall supporting the notion that cross-feeding plays an important role in pairwise interactions (Fig. 3c). To further explore metabolite exchanges, we constructed ecological networks from exometabolomic data, estimating metabolite production and consumption between strains (Fig. 3d, Interactive CrossFeed-o-Gram). This identifies metabolites predicted to mediate interactions. For example, net release of lysine, trehalose, sucrose, and cytidine by *Variovorax* and arginine, citric acid and alanine by *Paenibacillus* matches utilization by *Mucilaginibacter*, putatively explaining *Mucilaginibacter*’s high abundance in co-cultures despite a mediocre growth rate (Fig. 3d).

### From pairwise interactions to whole community dynamics

To examine interactions in a complex community, and test whether the CRM can recapitulate multi-species dynamics from monoculture parameters, we assembled complete 15-member communities in defined medium (NLDM) (Fig. 1d). Cultures were inoculated with equal OD and passaged every 48 hours for 10 days in two independent experiments (A, B), each with four replicates (Fig. 4a-b). Replicates were highly reproducible within experiments, but the two experiments stabilized at distinct compositions, potentially reflecting the effect of noise, stochasticity or multistability. Despite these differences, five strains coexisted at high abundance in both experiments: *Rhizobium, Mucilaginibacter, Lysobacter, Chitinophaga*, and *Burkholderia.* Three strains, *Arthrobacter, Variovorax* and *Niastella* additionally persisted in Experiment A (Fig. 4a). Further experimental insight into the sensitivity of these community compositions to perturbations can be obtained from leave-1-out and leave-2-out experiments (see below).

**Figure 4:**
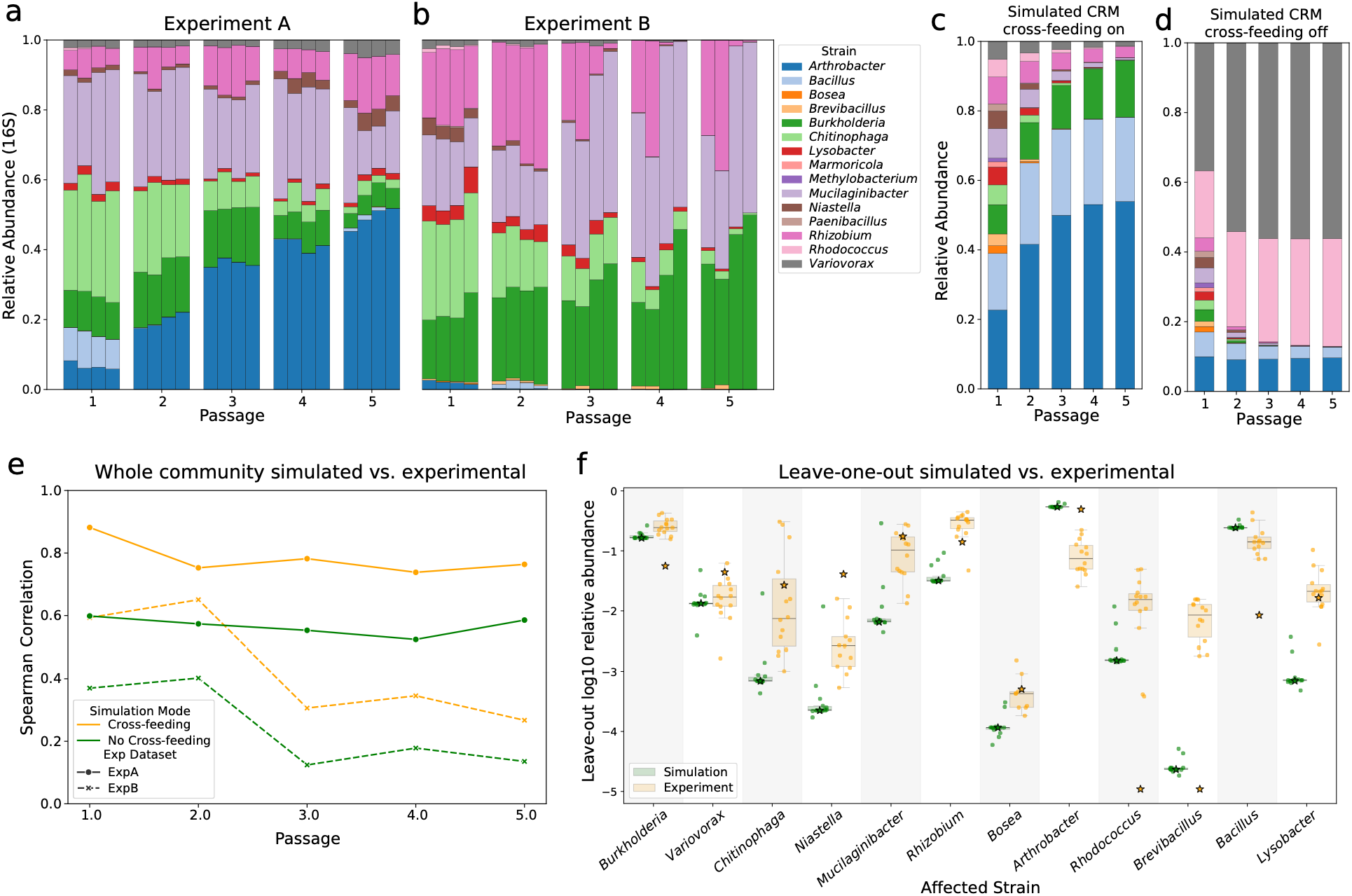
Experimentally-informed CRMs predict complex community assembly. a,b) Experimental dynamics of community composition in a synthetic ecosystem consisting of all 15 strains cultured for 10 days in 5 passages in NLDM media. 2 experiments were performed, each with 4 biological replicates. 16S relative abundances are shown as stacked bars. **c)** Whole community (15-member) simulated abundances across 5 48-hr periods with passages in between. Fitted CRM parameters are used, and relative abundances of strains in the final community are shown as stacked bars. **d)** Whole community (15-member) simulated abundances across 5 48-hr periods with passages in between. Cross-feeding was turned off by setting leakage parameters for every strain = 0 (Methods). Relative abundances of strains in the final community are shown as stacked bars. **e)** The correlation (Spearman’s coefficient) between the abundance ratio of 15 inoculated strains at each passage between average 16S measurements across 4 replicates and CRM simulations with cross-feeding (orange line) or CRM simulations without cross-feeding (green line). Strain abundances from simulations were compared to both instances of whole community experiments (dashed and solid lines). Spearman coefficients are used due to non-gaussian distribution of abundances. **f)** The simulated and experimentally observed strain abundances in the community across leave-out experiments of all other 14 members, one at a time. The y-axis shows the log abundance of the strain of interest across leave-out experiments of all other strains. The strain on the x axis indicates the strain of interest who’s abundance is plotted on y (e.g. Burkholderia’s relative abundance across all single leave-outs of the 14 other strains). The star for each represents that strains abundance in the complete 15-member community (accounting for the artificial removal of the left out strain so relative abundances are comparable). In green are the simulated predicted abundances, and in orange is the experimentally measured abundance.

We simulated the 15-member community with our monoculture-fitted CRM under the same passage scheme (Fig. 1e). The model predicted stable coexistence of six strains (Fig. 4c), three of which (*Burkholderia*, *Mucilaginibacter, Rhizobium*) dominated in all experiments. Model composition more closely matched Experiment A (r=0.76) than B (r=0.31), sharing 5 out of 6 dominant taxa with A (Fig. 4e). When metabolite exchange is artificially disabled (Fig. 4d), community structure is drastically disrupted, and agreement with experiments decreased substantially (A: ρ=0.58, B: ρ=0.11), with fast growers dominating (Fig. 4e). This underscores the importance of cross-feeding in community assembly and dynamics. These model-experiemnt correlations are robust to alternative normalizations of the 16S relative abundance data and to changes in the model’s initial conditions (Supplementary Fig. 9-11).

To determine how strains influence one another in community context and whether assembly is particularly sensitive to specific strains, we performed leave-one-out experiments both *in silico* and *in vitro* by systematically removing strains from the 15-member community and tracking assembly outcomes (Fig. 4f, Methods). One striking inference from *in vitro* data is that most strains either benefit from or are hindered by removal of any other member. For example, the proportion of *Niastella* decreases by an average factor of 10 when any other individual strain is removed, whereas *Burkholderia’s* abundance generally increases under the same perturbations. We found no clear relationship between pairwise coexistence patterns and leave-one-out effects in the whole community. This may reflect our particular synthetic community or a fundamental difference in how strains interact with each other as isolated pairs versus how they affect each other within the community, an effect shown by comparing the experimentally derived interaction matrices for the two cases (Supplementary Fig. 7).

As seen in Fig. 4f, the CRM predicts smaller community shifts upon single strain removals relative to experiments. Prediction accuracy was strongly strain-dependent, with effects on *Burkholderia* and *Variovorax* displaying best match to experimental data. When *Arthrobacter* was removed, the model correctly predicted high *Burkholderia* abundance, with an overall community composition resembling more closely Experiment B (ρ=0.56) than A (ρ=0.31) (Supplementary Fig. 12-13). This suggests that perturbations to the initial composition, specifically the presence of *Arthrobacter*, may bias the system toward distinct community states, consistent with divergence between Experiments A and B. Leave-one out tests also corroborated some coculture-inferred interactions. For example, *Mucilaginibacter,* predicted to cross-feed with *Variovorax* and *Paenibacillus*, performed much worse without these partners (Supplementary Fig. 12b). Together, the initial presence of key strains influences final community structure, highlighting the complexity of interactions in larger consortia and motivating a deeper analysis in the next section.

### Leave-two-out experiments reveal nonlinear strain interactions

From the previous experiments, we learned that removing specific bacteria strongly affects community composition. We next asked whether removing strain pairs could produce nonlinear effects. We thus developed a modified epistasis metric to quantify deviations from a simple additive model of species interactions (Methods). Inspired by prior formulations linking genetic modifications to fitness or species additions to community function ^48,49^, we propose a new metric that measures the nonlinear impact of jointly removing two strains on any other strain in the community. This way of quantifying interactions complements pairwise and leave-one-out metrics and provides additional insight into ecosystem dynamics.

Similar to genetic epistasis^50^, our metric captures overlapping or complementary functions between strain pairs, as detected through their impact on the rest of the community. Leave-one-out and leave-two-out removals were classified by direction (beneficial, deleterious, or reciprocal when removals have discordant direction) and compounding effect (synergistic or buffering). For example, a pair (*i*, *j*) exhibiting deleterious synergistic epistasis (ε_ijk_ < 0) on strain *k* indicates reduced fitness of *k* in both single-and double-leave-outs as compared to whole-community, with the leave-two-out fitness effect exceeding the expectation from single leave-outs (Supplementary Fig. 14). Conversely, buffering occurs when removing one individual species on top of another has a milder effect than predicted.

Epistasis values were calculated for all three species combinations (*i*, *j*, *k*), as the relative abundance of *k* after removing *i* and *j*, relative to null expectations from *i* and *j* single removal experiments. Our epistasis inference therefore constitutes a 3D tensor, similar to a previous multi-phenotype epistasis metric in metabolic networks ^50^. The epistatic distribution displayed the characteristic trimodal pattern ^48^, and most interactions were buffering regardless of direction (Fig. 5a). Most strains showed consistent interaction effect direction regardless of the strain pairs left out, potentially reflecting differences in invasiveness or self-sufficiency (Supplementary Fig. 15) also seen in single leave-out experiments (Fig. 4f). Some strains, like *Arthrobacter,* experienced both buffering and synergistic epistasis dependent on the pair removed (Fig. 5b,5d), whereas others, like *Burkholderia* and *Niastella,* experienced primarily buffering epistasis regardless (Fig. 5c). The very different way in which *Arthrobacter* and *Burkholderia* respond to perturbations of other species may help explain the different whole community compositions observed in Experiment A and B (Fig. 4a-b), where *Arthrobacter* and *Burkholderia* were correspondingly dominating at the expense of the other.

**Figure 5.**
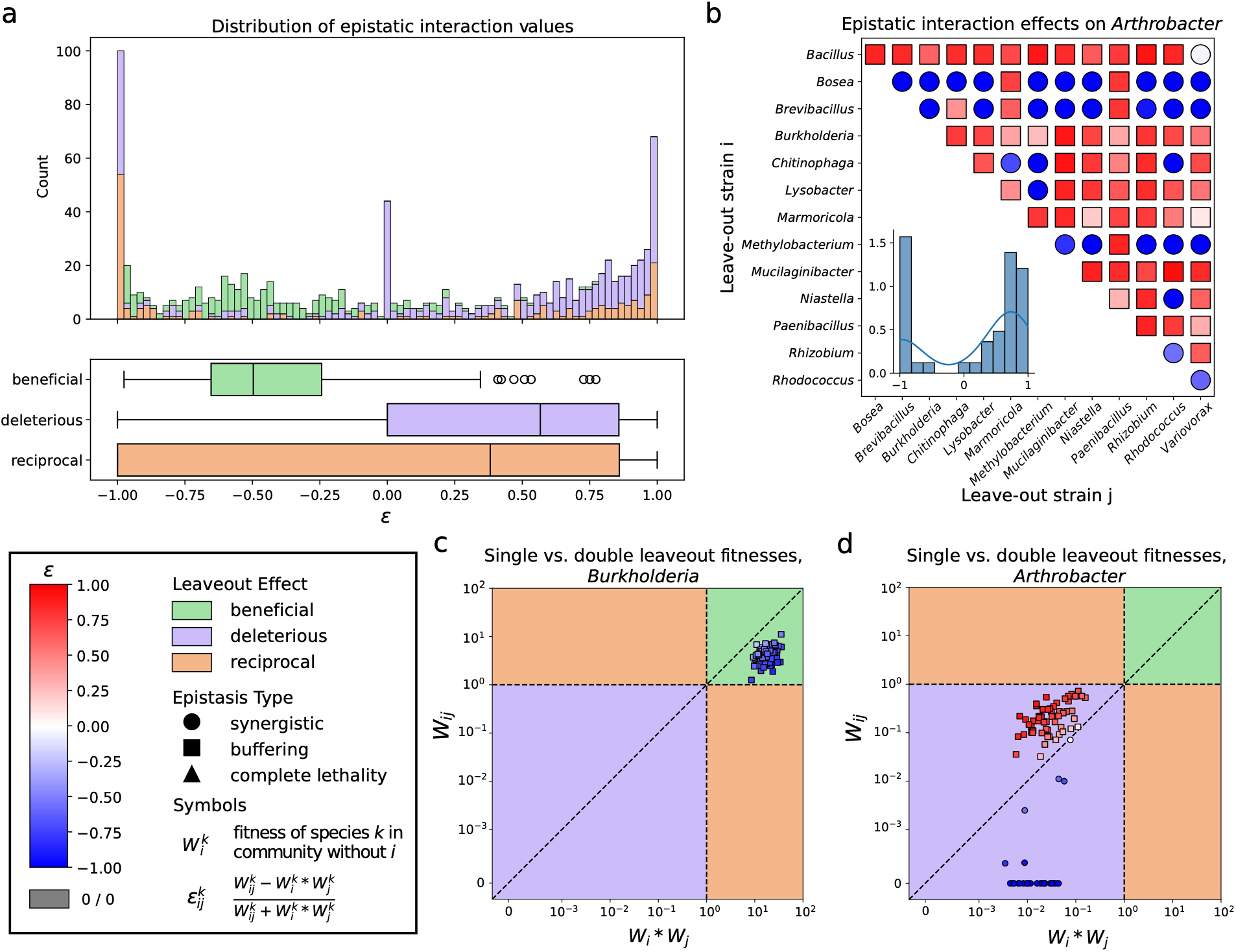
Epistatic interactions are prevalent across synthetic community members. **a**) Distribution of 910 epistasis values calculated from each trio of the 15 species, excluding 5 species from effect consideration due to little growth in whole-community. Peaks are observed at-1 (complete lethality), 1 (complete recovery), and 0 (no epistatic effect). Interactions are colored by direction of the leave-out effect on the target species. Most interactions are buffering (beneficial and *ε* < 0, or deleterious and *∈* > 0). See Supplementary Fig. 14 for more detail on epistasis and examples of classifying community interactions. **b)** Pairwise epistatic interaction effects on *Arthrobacter* relative abundance. Some combinations result in synergistic epistasis (circles) while others result in buffering epistasis (squares). Each pair is colored by the epistasis value. All leave-outs exhibit deleterious effects, as seen in D. **c)** Comparison of double leave-out effect on *Burkholderia* fitness against expected fitness from linear combination of single leave-out effects. All pairs exhibit buffering beneficial effects, indicated by squares in the green region. **d)** Same as C, for *Arthrobacter*. Some pairs exhibit buffering epistasis (squares), while some exhibit synergistic epistasis (circles). All pairs exhibit deleterious effects, indicated by the purple region.

Notably, leave-out experiments also pointed to extreme cases where leave-out pairs caused complete lethality of target species that had increased fitness in both single leave-outs, while other pairs induced recovery of target species which would not grow in leave-one-out cases (Supplementary Fig. 15). This lethality, similar to synthetic lethality in genetic epistatic networks, may point to redundancy of functions ^48,51^.

### Model-driven design of diversity-enriched subcommunities

Beyond validating predictions against existing community assemblies, we next asked whether our CRM could be used prospectively to design and test entirely new communities selected for a target ecosystem-level property. The real value of the model lies in creating large in silico community sets, which would be impractical to construct experimentally, and ranking them by their predicted likelihood of displaying a desired behavior.

Because species diversity influences community resilience, ecosystem-level functions, and host health ^52,53^, we used the CRM to identify synthetic subcommunities with enriched species diversity. We simulated the assembly of all possible (N=455) three-species communities over five passages and calculated the Shannon entropy for all final compositions (Fig. 6a). From these, we selected 10 communities predicted to display coexistence, high OD, and high Shannon entropy, indicating balanced communities. As controls, we chose 10 communities ranked lowest in Shannon entropy, while still maintaining coexistence. These 20 communities, predicted to rank at the top and at the bottom of our desired community-level phenotype, were assembled experimentally, and passaged for 10 days every 48 hours, and profiled by 16S sequencing (Fig. 6b).

**Figure 6.**
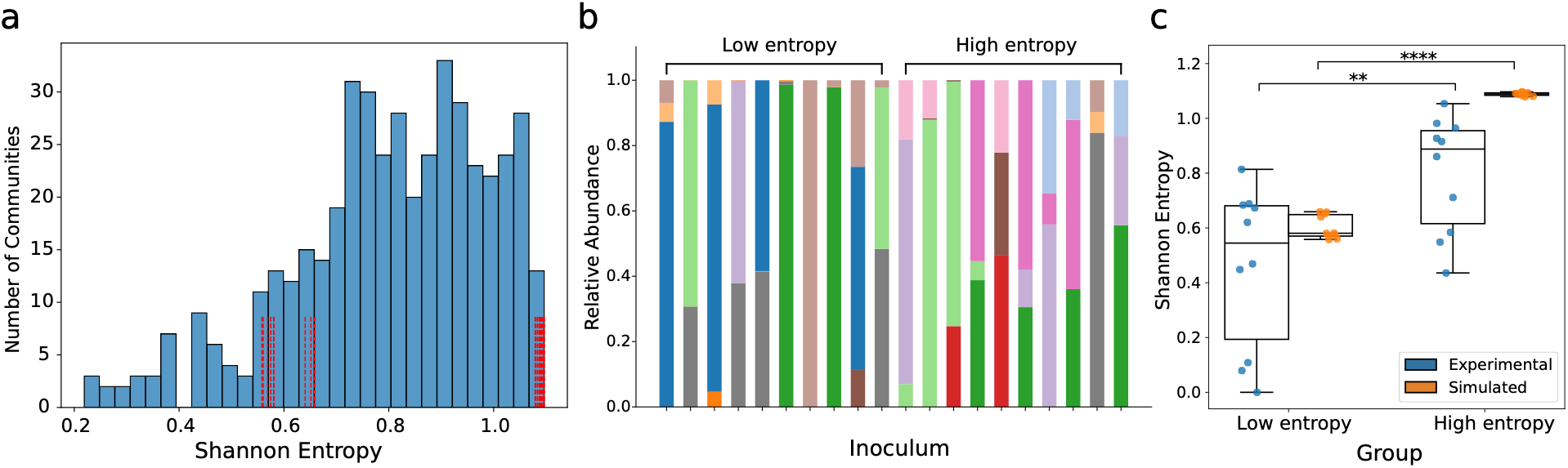
CRMs generate synthetic sub-communities with high species diversity. **a)** Distribution of the Shannon entropy of 455 3-strain synthetic sub-communities assembled *in silico*. The red vertical lines specify the communities chosen for the experiments shown in Fig. 6b. Red lines at the right end of the distribution correspond to communities in the high predicted Shannon entropy group, while red lines towards the lower end of the distribution correspond to those in the low predicted Shannon entropy group. **b)** The taxonomic composition of selected groups of 3-strain subcommunities (Supplementary Table 4) at the end of a 10-day growth experiment. There are 10 sub-communities containing 3 strains in the initial inoculum predicted to have a low Shannon entropy (<0.7) on the left compared with 10 sub-communities predicted to have a high Shannon entropy (>1). **c)** Distribution of the Shannon entropy of selected 3-strain sub-communities from the 16S measurements (“Experimental”) and from simulation (“Simulated”) at the end of the final passage (Day 10). Two-sample two-sided independent t-tests (df=18) were performed for the statistical significance between groups for the 16S measurements (p<0.01) and the simulations (p<0.0001).

Top ranking communities displayed significantly higher experimental Shannon entropy than controls (p = 0.008). Although experimental diversity was more variable and generally lower than the corresponding predictions, the modeling and experiments agreed in overall trends and group differences (Fig. 6c). These results demonstrate that experimentally parametrized CRMs can inform the design of communities with desired ecosystem-level properties by screening large in silico community sets and narrowing the search space for experimental validation and community engineering.

## Discussion

Among dynamical models of microbial community assembly, consumer-resource models (CRMs) fit between the high detail of reaction stoichiometry in genome-scale metabolic models (GEMs) and the coarse interaction structure of generalized Lotka-Volterra models. CRMs have been most widely used to study principles of coexistence, stability and species richness by sampling parameters from distributions ^39,40,43^ or inferring them based on structural properties of real metabolic networks ^20,42,45^. Our study adds to a small but growing effort to use CRMs as predictive tools parametrized with experimental metabolomics data and applied to a specific model system ^28,54^. One previous work used untargeted metabolomics to build a CRM with coarse metabolite groups as resources, finding that resource competition rather than cross-feeding determines structure in gut-associated communities ^28^. Another study parametrized a minimal two-resource CRM of denitrification across dozens of soil isolates, predicting community metabolite dynamics from monoculture phenotypes alone ^54^. Within this landscape, our work adds several novel elements. Our rhizosphere CRM is based on targeted exometabolomics data that enable parametrization of the model for individual metabolites known to be relevant plant exudates. This rich metabolic phenotyping data allows us to trace organism-specific uptake and secretion patterns mediating competition and cross-feeding. It is also worth noting that our effort focused on a set of microbes that is becoming a standardized synthetic community among plant microbiome researchers ^25,55–58^. We anticipate that this type of convergence of multiple labs and measurements on defined synthetic communities will deepen understanding of microbial ecosystem processes ^59^. Finally, in addition to enabling the inference of mechanistically interpretable microbial interaction networks, our work shows a viable approach to estimate the community-level effects of specific interventions (e.g. removal of a metabolite or a consumer) and design de-novo communities with desired properties (Fig. 6).

By inferring from metabolomics data a D matrix (internal transformation across metabolites) specific to each strain, we capture distinct metabolic strategies that would be absent in most CRMs, which typically assume the same D matrix across all species. The D matrix determines exudation capabilities of each organism, and is therefore instrumental for predicting cross-feeding. Notably, when cross-feeding is artificially removed from the model, letting competition for resources determine the dynamics and enabling fast-growing strains to dominate, predictive accuracy is reduced (Fig. 4d). This suggests that cross-feeding is necessary to capture dynamics in systems where composition is not primarily explained by differences in intrinsic growth rates. This is consistent with the finding that obligate cross-feeding interactions appear to be a general feature of soil communities rather than an exception ^60^. Modeling metabolite exchange at this resolution does, however come at a cost. Fitting strain-specific metabolite exudation for 64 resources amounts to 4096 parameters per strain (∼85% of which are 0), which we estimate using a combination of direct analytical inference from exometabolomics and subsequent optimization. This carries an obvious risk of overfitting: current parameters are optimized against a specific set of monoculture data and would likely need to be modified for growth in a different nutrient context.

While our study suggests that both cross-feeding and competition constitute important metabolite-mediated interactions, the comparison of different experiments indicates that these interactions may look different in pairwise versus community context. For example, while *Burkholderia* does not show a strong direct interaction with *Arthrobacter* in the pairwise co-culture (Fig. 2a), the two organisms seem to be in conflict to dominate in the whole community context (Fig 4a, b). This phenomenon is observable in the whole community repeat experiment (with each of the two organisms dominating in one of the repeats) and is consistent with the leave-out-experiments (Fig. 4f), as well as with recent reports of *Burkholderia* dominating similar communities ^25^. Metabolite-mediated interactions in our experiments may be strongly modulated by the molecular composition of the medium. For example, *Methylobacterium*, whose preferred carbon source (methanol) is not present, may be secreting unusable byproducts of non-ideal carbon sources ^61^. For other strains, secretion may be a result of overflow metabolism ^62^, or a greedy usage of resources that result in stoichiometric elemental imbalance and advantageous exudation of unusable moieties ^63^. In general, our different types of interaction matrices (pairwise, leave-1-out, epistasis) could be viewed as complementary descriptors of how the bacteria interact in different contexts. No single definition captures all interaction layers, supporting the need for multidimensional metrics ^64^.

Two measurement limitations bound our conclusions, and each suggests a concrete improvement for future iterations of this approach. First, experimental and simulated compositions are not quantified on the same footing. On the experimental side, we report 16S rRNA relative abundances, which are subject to well-documented technical biases due to differences in copy number, extraction efficiency, and amplification ^65–67^. Conversely, our model is parametrized from monoculture optical density and predicts biomasses, which are then converted to relative abundances. Given the multiple possible sources of uncertainty, we show that community results are stable across alternative 16S normalizations (copy number, and colony forming unit to OD conversions), and varying model initial conditions (See systematic analysis in Supplementary Methods 2, Supplementary Fig. 9-11). Direct comparisons at the level of individual abundances (beyond the rank-based metrics we used) would require absolute calibration, with spike-in standards, qPCR, or cell counts, which we would recommend building in from the outset in future work. Second, our metabolomics analysis quantified only compounds present in NLDM. Although NLDM is metabolically rich, unmeasured metabolites, especially bacterially-produced metabolites missing from the starting media, likely play important roles as exchanged nutrients or signals ^68,69^. For instance, a metagenomic-based network analysis has recently shown that thiamine released by the same *Variovorax* in our collection is an essential vitamin that can support auxotrophic partners ^55^. Consistent with this, several leave-one-out perturbations had strong experimental effects missed by the model, suggesting the existence of yet unobserved influential metabolites or indirect metabolic dependencies. It is also possible that untracked carbon-containing metabolites significantly affect the growth of some strains; for example, *Arthrobacter* grows to a final yield lower than expected based on total carbon uptake suggesting exudation of untracked metabolites (Supplementary Fig. 3c). In future work, this limitation could be addressed through additional untargeted metabolomics, which would widen chemical coverage.

Beyond measurement limitations, several biological processes are not accurately captured by our CRM. The D matrix represents internal resource transformations in a coarse manner, while real metabolic fluxes depend on carbon/nitrogen balance, oxygen, pathway architecture, and diauxic shifts. Moreover, both the C and D matrices are fixed over the course of a simulation, so time-dependent shifts in metabolic strategy, such as sequential substrate use, cannot be represented. Although aspects of this may be implicitly captured in the learned parameters through simulated annealing, many properties of metabolism are likely oversimplified or omitted, a limitation that could be addressed using stoichiometric or dynamic flux balance models. Comparing CRM performance against GEMs available for these 15 species ^56^ would in fact be a very informative future endeavor. Although our CRM achieved reasonable accuracy, next iterations of our approach would greatly benefit from improved and more flexible parameter fitting.

Looking forward, our findings highlight both opportunities and challenges for microbiome engineering. This framework could guide efforts to engineer environments to enrich taxa or steer communities toward alternative states. While we focused on taxonomic composition and diversity prediction, similar approaches could target functional objectives, such as stabilizing metabolic activities or suppressing pathogens, as shown for *C. difficile* ^70,71^. The chemical diversity of NLDM creates many niches, making collective functions difficult to predict, yet offering a rich testbed for linking metabolic diversity to community traits. Finally, bridging model simplicity and biological realism remains crucial. Alongside mechanistic models, growing work highlights the potential of data-driven ML/AI-based approaches to predict community assembly and function, offering complementary strengths when mechanistic detail is incomplete ^72,73^. Continuing to develop intermediate frameworks between CRMs and GEMs and combining mechanistic models with data-driven frameworks could reveal how coarse-graining constrains both predictive power and design potential in microbial communities.

## Methods

### Growth medium

We used Northen Lab Defined Medium (NLDM) for all monoculture, co-culture, whole community and sub-community experiments, which is a defined medium specifically designed for cultivation and exometabolomics profiling of soil bacteria^27^. Metabolites included and concentrations are found in Supplementary Table 2 and more information on the specific media recipes can be found on protocols.io ^74^. Metabolites were included based on metabolites found in rhizosphere environments and metabolites in a popular undefined media for cultivating soil microbes, R2A. Metabolites in NLDM were categorized throughout the manuscript in 5 classes (sugars, amino acids, organic acids, nucleotides/nucleosides and others) and are reported in Supplementary Table 2.

### Selection and reviving of strains

The model synthetic soil community obtained from the rhizosphere and surrounding soil of a switchgrass include 15 bacterial isolates (Supplementary Table 1) (*Lysobacter* spp. OAE881, *Burkholderia* spp. OAS925, *Variovorax* spp. OAS795, *Chitinophaga* spp. OAE865, *Niastella* spp. OAS944, *Mucilaginibacter* spp. OAE612, *Rhizobium* spp. OAE497, *Bosea* spp. OAE506, *Methylobacterium* spp. OAE515, *Arthrobacter* spp. OAP107, *Rhodococcus* spp. OAS809, *Marmoricola* spp. OAE513, *Brevibacillus* spp. OAP136, *Paenibacillus* spp. OAE614, *Bacillus* spp. OAE603) ^2^. Each organism was recovered on 1X R2A agar (Teknova, CA) from its glycerol stock, and stored at 4 °C for no more than 30 days. Single colonies from individual isolates were grown in 2 mL liquid cultures of Northern Lab Defined Media (NLDM) at 30 °C with shaking at 200 rpm for 72 hours.

### Culturing of microbial monocultures and communities

For monoculture and following experiments, the starting culture of selected 15 bacterial organisms were grown separately in 3 batches. Single colonies of *Methylobacterium* and *Marmoricola* spp. were cultured in 2 mL NLDM (x5) for 72 hours, and colonies of *Bosea*, *Brevibacillus*, *Paenibacillus* and *Bacillus* spp. were grown in 2 mL R2A (x3) for 48 hours prior to pooling. The starting cultures of remaining isolates were grown in 2 mL R2A overnight. All cultures were incubated at 30 °C with shaking at 200 rpm. Multiple inoculums from the same isolate were pooled to obtain enough biomass for the subsequent growth assay. Each cell culture was washed two times in PBS followed by NLDM, and pelleted by centrifugation at 4,000 x g for 5 min and removing the supernatant. Collected pellets were resuspended in 100 µL NLDM, and dilution of cell resuspension at 1:20 was measured for OD_600_. OD_600_ of each culture was adjusted to be 0.15 for further dilution.

To assay for monoculture growth, each cell culture was diluted in 200 µL NDLM with an initial OD600 0.1. Two technical replicates of each isolate except *Methylobacterium* and *Marmoricola* spp. were grown at 30 °C and 286 rpm in flat bottom 96-well plates for 72 hours. Parallelly, two replicates of each culture were incubated in the microplate reader at 30 °C with continuous orbital shaking at 286 rpm. Values of OD_600_ were read every 20 min for cultures grown in the microplate reader. When the OD_600_ values of replicates from each isolate in the microplate reader exceeded 1.0, cultures from the selected isolate grown in the incubator were sampled periodically for OD_600_ reading after appropriate dilution spanning its exponential growth phase. The growth profiles of the 13 organisms grown in an incubator, and *Methylobacterium* and *Marmoricola* spp. grown in the microplate reader are shown in Supplementary Fig. 2.

### Growth of coculture pairs and higher order communities

For experiments involving co-culturing of 2 or more organisms, cultures were diluted and combined in biological replicates in a 200 uL final volume with an initial OD_600_ 0.01 of each organism. Cocultures were incubated at 30 °C and 286 rpm in flat bottom 96-well plates for 48 hours and were further passaged 2 times. For each passage, 5 uL of the grown culture was transferred into 195 uL fresh NLDM medium. For experiments involving the whole or a subset of the 15 organisms, cultures were combined in biological replicates at final concentrations of OD_600_ 0.01 in 500 uL NLDM, and passaged every 48 hours for a total of 10 days. At each passaging step, 10 uL of the grown culture was transferred into fresh medium for the subsequent growth period, 10 uL was transferred into 190 uL PBS for OD_600_ measurements.

Communities to be sequenced were centrifuged at 4000 x g for 10 min and the supernatant was removed. Cell pellets were stored at-80 °C before use. DNA was collected using a 96-well genomic DNA purification kit (Invitrogen), and the manufactures protocol was performed.

### Amplicon sequencing

The 16S ribosomal RNA gene amplicon libraries were prepared for sequencing on the Illumina MiSeq platform at Quintara Bio, Boston, MA. The V4 region of the 16S rRNA gene was amplified with the primers U515F (5’-GTGCCAGCMGCCGCGGTAA) and E786R (5’-GGACTACHVGGGTWTCTAAT), developed by Preheim et el. ^75^. All raw 16S sequencing data for each study was separately processed using BU16S (https://github.com/Boston-University-Microbiome-Initiative/BU16s), a QIIME2 ^76^ pipeline customized to run on Boston University’s Shared Computing Cluster. Briefly, BU16S first trims primers and filters out reads of less than 50 base pairs using cutadapt, then obtains ASVs using dada2 ^77^, and finally classifies ASVs with 95% or greater sequence identity to the SILVA_132_99 database with VSEARCH ^78^. Compositional vectors for different communities were computed by diving reads for individual strains by the total number of reads. After determining that there was no obvious advantage to choosing specific normalization strategies, and that our main results are robust to these different possible corrections (see Supplementary Methods 2, Supplementary Fig. 9-11), we decided to report the results with no correction factors.

### Exometabolomics Sample Preparation

Quadruplicate 1.2 mL cultures (including medium controls) of the selected 15 bacterial organisms were cultured in NLDM at 30°C. Multiple time points were collected from individual wells, at 4, 5.5, 8.5, 10, 12.5, 19.5, 29, 29.5 and 44.5 hours, roughly matching with 25%, 50%, 75% and 100% growth yields of each strain. A culture fraction of 1 mL was centrifuged at 7,000 x g for 5 min at 4 °C and 0.75 mL of the supernatant was collected. For *Methylobacterium*, *Marmoricola* and *Paenibacillus* spp., only the timepoint corresponding to the maximum yield was sampled due to challenges in collecting intermediate growth stages. The supernatants were then frozen at −80 °C, lyophilized and resuspended in 375 μL methanol containing internal standards ^79^. The resuspended samples were filtered through 0.2μm modified nylon membrane centrifugal filters (Pall Corporation, Port Washington, NY, USA) for 2 min at 5,000 x g and analyzed as described in the LC-MS/MS section.

### LC-MS/MS analysis and metabolite identification and statistical analysis

Metabolites in the NLDM were chromatographically separated using hydrophilic interaction liquid chromatography (HILIC) and detected by high resolution tandem mass spectrometry ^79^. Analyses were performed using an InfinityLab Poroshell 120 HILIC-Z column (Agilent, Santa Clara, CA, USA) on an Agilent 1290 stack connected to a Q-Exactive Hybrid Quadrupole-Orbitrap Mass Spectrometer (Thermo Fisher Scientific, Waltham, MA, USA) equipped with a Heated Electrospray Ionization (HESI-II) source probe. Briefly, metabolites were separated by gradient elution followed by MS1 and data dependent (top 2 most abundant MS1 ions not previously fragmented in last 7 s) MS2 collection; targeted data analysis was performed by comparison of sample peaks to a library of analytical standards analyzed under the same conditions. Three parameters were compared: matching m/z, retention time and fragmentation spectra using Metabolite Atlas (https://github.com/biorack/metatlas). Metabolite background signals detected in the extraction blanks were subtracted from the experimental sample peak heights/areas. Metabolite peak heights and peak areas were normalized by setting the maximum peak height or peak area detected in NLDM uninoculated control samples to 100%. Normalized metabolite feature peak heights or peak areas were used for relative abundance comparisons.

### Consumer-resource model

To simulate microbial population dynamics of the growth of single species, or arbitrary subsets of the synthetic community, we employed an ODE-based microbe-metabolite modeling approach based on the classic MacArthur consumer resource model ^80^ which was modified to incorporate resource exchange and preference in a microbial community ^39,81^. In this modeling approach, the set of ordinary differential equations below governs the dynamical change of population abundances *N_i_* of S microorganisms, and resource abundances *R_α_* of M metabolites. A detailed walkthrough of these equations and how each term is biologically relevant can be found in the Supplementary Methods 1.1.

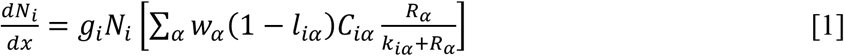

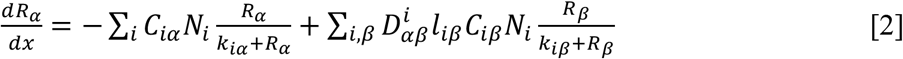

The population abundances *N_i_* (i = 1,2,…,S) are determined by (i) a stoichiometric resource utilization matrix *C_iα_*, (ii) a species-specific conversion factor from energy uptake to growth rate *g_i_*, and (iii) a scaling factor for energy content of resource α, w. The uptake rate per unit concentration of resource α by organism I, as captured by Ciα multiplied by the availability of resource α with a Monod kinetics parameter 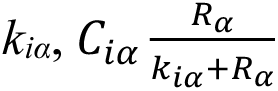 models the growth of each microbial species on each resource. After uptake, proportions of resources *β*, *l_iβ_*, are converted to resources α, encoded by a species-specific normalized stoichiometric D matrix *D*_α_*_β_^i^*, and secreted into the environment as common goods for the rest of the community organisms. The remaining fraction of resources α, 1-*l_iα_*, contributes to the growth of each organism. The dynamic variables and mechanistic parameters are defined in Supplementary Table 3, and a more in-depth explanation of the model is present in the supplement (see Supplementary Methods 1.1).

### Initial parameter estimation for CRM

We estimated the parameters in the resource utilization matrix *C_iα_* and the growth rate conversion factors *g_i_* from energy uptake, using time-course measurements of exometabolomics, detailed in the Supplementary Methods 1.2. The kinetic growth curve of *Lysobacter* was used to estimate the orders of magnitude of the remaining parameters. The energy content of each resource w was set to 1 x 10^12^ and the Monod resource consumption half-velocity constant was set to 0.04 g/mL for all resources to approximate the timeframe of growth. The minimum energy requirements for all organisms were assumed to be 0. Lastly, the leakage fraction lα was assumed to be species-dependent and set to 0.2 for all resources in each species.

### Simulated annealing optimization of CRM parameters

After obtaining an initial estimation of parameters used in consumer resource modeling, we implemented a Monte Carlo simulated annealing algorithm to tune the set of parameters to minimize the population and metabolite usage differences between simulated monoculture growth of each species and the experimental measurements. The search initializes with the parameters quantified above. Initial species abundance was set to 1 x 10^7^ CFU/ml. Each iteration makes perturbations to parameters in *C_iα_* and *D_i_*, as well as *g_i_* and *l_iα_*. A cost function

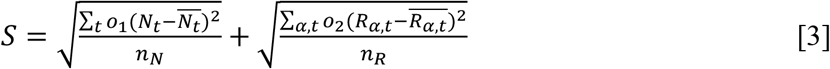

Was used to evaluate the performance of parameters at each iteration, in which *N_t_* ^_^ and *R_α_*_,*t*_,^_^ t are the simulated species population abundance and resource abundance at each sampled timepoint *t*, whereas *N_t_* and *R_α,t_* are the population abundance determined from experimentally measured OD_600_ and resource abundance from the exometabolite measurement. The score represents the distance between simulation and experimental measurements, while the contributions of population abundance and resource abundance are weighted differently according to the *o*_1_ and *o*_2_ factors. Details of the simulated annealing algorithm are provided in the Supplementary Methods 1.3. Simulation of monoculture growth from the improved parameter sets is shown in Fig. 2c, and the respective resource depletion or secretion is shown in Fig. 2d.

### Implementation of CRM simulations for community dynamics

The ODE solver used for all CRM simulations was scipy.integrate.odeint from scipy v1.14.0 with parameter mxstep set to 50 and timesteps set to 0.01 (∼36 seconds). Initial population abundances in simulated experiments were consistently set to 0.01 per species (to reflect experimental OD initial conditions), and then scaled by a constant OD-to-CFU conversion of 10^9^. Simulated experiments were performed to replicate their experimental counterpart in terms of length of time and number of passages. Passages were performed such that the final species abundances at the end of one passage were scaled by 0.02 and assigned as the starting abundance of the next passage, mimicking the same dilution factor as the in vitro experiments. Resource abundances for all experiments reflect the media composition of NLDM in units g/mL.

### Interactions from Consumer-Resource Model Parameters

Using the experimentally-informed consumer resource models with parameters fitted and optimized, we derived species-species and species-metabolite interactions based on the set of *C_iα_*, *D_i_* and *l_iα_* parameters. Here, we define the interaction strength between species *I* and metabolite *α* as a summation of its positive interaction strength (*e_αi_*), the rate (mL/hr) at which resource α is generated by species *I*, and its negative interaction strength (*e_iα_*), the consumption rate (mL/hr) of resource α by species *i*. The derivation of *e_αi_* and *e_iα_* is as follows:

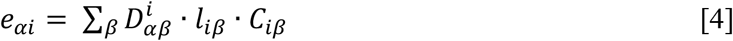

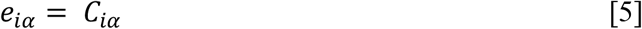

### Construction and visualization of ecological networks

Ecological networks were constructed using the Networkx v 2.6.3 and visualized using PyVis v 0.3.2. Metabolite sharing between species is inferred by the production and consumption values of exometabolites, with large production values constrained for ease of interpretation of edge weights (see ‘LC-MS/MS analysis and metabolite identification’). Representative networks were constructed by creating the whole community network, then creating a sub-graph of the species of interest. Dead-end nodes (production or consumption of a metabolite that is not produced or consumed by other species represented in the network) are trimmed. The jupyter notebook “CrossFeed-o-Gram” can be used to explore putative interactions between species.

### Interactions and epistasis values from leave-out experiments

Relative abundances were calculated for the whole-community, leave-one-out, and leave-two-out experiments and averaged across replicates. Fitness *W_X_^k^* of each species *k* under leave-out *X* was calculated by comparing relative abundances ω*_X_* against whole-community abundances ω_W*C*_ and scaling based on left-out species:

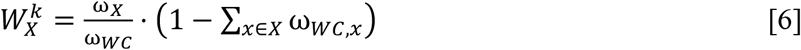

By this formula, whole-community fitness is 1 for all and thus ignored in our epistasis calculation. Because fitness values can range from 0 to large positive values, a normalization factor was applied to the traditional epistasis formula to constrict values to the range [-1, 1]. Epistasis is calculated as

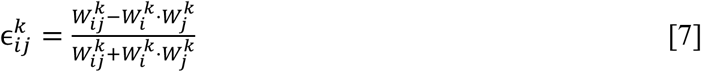

where *ɛ_ij_^k^* quantifies the interaction between leave-out species *i* and *j* in terms of its effect on species *k*. Interactions were classified into three groups based on effect direction: beneficial (*W_ij_^k^*> 1 *and W_i_^k^* ⋅ *W_j_^k^*> 1), deleterious (*W_ij_^k^* < 1 *and W_i_^k^* ⋅ *W_j_^k^* < 1), and reciprocal (*W_ij_^k^* < 1 *and W_i_^k^* ⋅ *W_j_^k^*> 1 or vice versa) (Supplementary Fig. 14).

### Prediction and testing of synthetic sub-communities

All possible 455 combinations of 3-species from the 15 species were simulated. The Shannon entropy as a measure of the final diversity after the 7 passages in the community was calculated in the standard way. We ranked the 455 3-species communities following an order of these metrics: (i) we considered only communities with co-existing (viable) species with a population number (biomass) higher than the equivalent of OD_600_ 0.1, (ii) we did not include communities with the presence of the slow-growing *Marmoricola* or *Methylobacterium*, (iii) we ranked them according to their Shannon entropy of the community, and (iv) if two communities had similar entropy, we ranked them by the community fitness reflected by the total population abundance of the community. From this ranking, we selected the top 10 communities and the last 10 communities (Supplementary Table 4). Ten of the 20 chosen communities were ones that resulted in relatively low diversity and 10 were with high diversity at the end of the simulations. Experiments for these 20 selected subcommunities were performed as described above and Shannon entropy was calculated from final 16S community.

## Data Availability

LC-MS data generated as part of this study are available from MassIVE under accession number MSV000103252. All raw 16S sequencing data generated in this study has been deposited in the NCBI Sequence Read Archive (SRA) under BioProject accession number PRJNA1531197. All data tables and code used for this manuscript is available at https://github.com/segrelab/metaCRM_SynCom.

## Authors Contributions

DSeg, JZ, TN and MdR designed the initial study; JZ, MdR, KH, and MO carried the experiments; SB, JZ, RN, DSco, DSeg, ID implemented computer models and analyzed the data; SB, JZ, RN, DSeg wrote the manuscript. All authors edited and approved the final version of the manuscript.

## Acknowledgements

This work was supported by the U.S. Department of Energy, Office of Science, Office of Biological & Environmental Research through the Microbial Community Analysis and Functional Evaluation in Soils SFA Program (m-CAFEs) under contract number DE-AC02-05CH11231 to Lawrence Berkeley National Laboratory; the National Science Foundation (grants NSFOCE-BSF 1635070 and the NSF Center for Chemical Currencies of a Microbial Planet-C-CoMP article #085); and the Human Frontiers Science Program (RGP0060/2021). Additional support was provided by the NIH (T32GM150533, T32GM100842), the NSF Research Traineeship at Boston University (DGE 1735087), and the Boston University Rafik B. Hariri Institute Graduate Student Fellowship in Computing and Computational Science & Engineering.

## Competing Interests

The authors declare no competing interests.

## Notes

### Competing Interest Statement

The authors have declared no competing interest.

### Summary of Updates

The manuscript was updated based on reviewer's feedback. Two additional authors were added reflecting their contribution to the revised manuscript.

