## Supplementary Material for "Metabolic blueprints of monocultures enable prediction and design of synthetic microbial consortia"

### Supplementary Methods

#### 1 Consumer resource modeling

##### 1.1 Overview of the model

Consumer-resource models describe microbial communities by keeping track of two sets of variables: the abundances of all the microbial strains or species (consumer $i$) and the environmental concentrations of each molecule or metabolite (resource $\alpha$). The main idea is that interactions between species are not imposed a priori but rather emerge from the propensity of individual consumers to use resources from (or contribute resources to) a shared pool of metabolites. This approach enables competition and cross-feeding to both be emergent properties of the metabolic utilization and exudation preferences of each organism. First, if two species consume the same metabolite, each depletes what is available to the other, and the strength of that competition follows from how much their consumption rates overlap across metabolites rather than from imposed competition coefficient. Second, in terms of cooperation, species may not use for growth or maintenance everything they uptake: the fraction they release back into the environment as byproducts becomes a resource that another species may be able to grow on.

The model consists of a set of coupled differential equations, one for metabolite abundances (Equation [S1]) and one for species populations (Equation [S2]). Supplementary Table 3 describes in detail the units of each variable in these equations and whether its value varies between strains and metabolites or is fixed.

$$\overbrace{\frac{dR_\alpha}{dt}}^{\text{Change in } R_\alpha} = \overbrace{\underbrace{-\sum_i C_{i\alpha}\, N_i}_{\text{Sum of sps. cons. of } R_\alpha}\ \underbrace{\frac{R_\alpha}{k_{i\alpha}+R_\alpha}}_{\text{kinetic term}}}^{\text{Metabolite } R_\alpha \text{ consumed}} + \overbrace{\underbrace{\sum_{i,\beta} D^i_{\alpha\beta}\, l_{i\beta} C_{i\beta} N_i}_{\text{Sum of sps. Prod. Of } R_\alpha \text{ from } R_\beta}\ \underbrace{\frac{R_\beta}{k_{i\beta}+R_\beta}}_{\text{kinetic term}}}^{\text{Metabolite } R_\alpha \text{ produced}} \qquad [S1]$$

The resource equation [S1] tracks how each metabolite's concentration changes over time. The first term represents consumption by the different species and has therefore a negative sign. This consumption term is calculated by summing over all species $i$ and multiplying individual consumption preferences for resource $R_\alpha$ by the size of the population $N_i$. The production term is positive because it returns metabolites to the shared pool, and it is built up of three components.

First, $C_{i\beta}N_i \cdot R_\beta/(k_{i\beta} + R_\beta)$ is the rate at which species $i$ takes up some other metabolite $R_\beta$. Second, multiplying by the leakage parameter $l_{i\beta}$, only a fraction of what was consumed gets released back into the environment. Third, multiplying by $D^i_{\alpha\beta}$ selects the set of other metabolites that end up releasing our metabolite of interest, $\alpha$ ,when they are consumed. $D^i$ is therefore a square matrix that represents a map of the internal metabolic capabilities of species $i$, determining the consumer-specific propensity that consumption of a given metabolite will lead to exudation of another metabolite into the environment.

$$\overbrace{\frac{dN_i}{dt}}^{\text{Change in sps. } i} = \overbrace{g_i N_i}^{\text{growth term and initial pop.}} \left[\sum_\alpha \underbrace{w_\alpha}_{\text{energy term}} \overbrace{(1 - l_{i\alpha})C_{i\alpha}}^{\text{proportion of } R_\alpha \text{ kept by sps } i} \underbrace{\frac{R_\alpha}{k_{i\alpha}+R_\alpha}}_{\text{kinetic term}}\right] \quad \text{[S2]}$$

The consumer equation [S2] reflects the fact that a species grows in proportion to the resources it retains from its environment, in the form of metabolite uptake. Growth of each consumer is also proportional to the consumer population itself ($N_i$), and to a coefficient $g_i$ that converts up taken metabolites into units of biomass in CFU/mL for that species. In the bracketed term, the Monod-like kinetic term $R_\alpha/(k_{i\alpha} + R_\alpha)$, multiplied by $C_{i\alpha}$, determines how fast one unit of biomass of species $i$ takes up metabolite $\alpha$. The factor $(1 - l_{i\alpha})$ imposes that a consumer- and resource-specific fraction $l_{i\alpha}$ is lost due to leakage/exudation into other metabolites. The term $w_\alpha$, originally introduced in CRMs to capture the energy content of metabolite alpha, can be thought more broadly as quantifying the contribution of metabolite $R_\alpha$ to growth.

In this version of the CRM, there are no terms for maintenance, death, or dilution, partially reflecting the fact that our experiments were performed as batch cultures. This model, as used throughout the paper, rests on additional simplifying assumptions. In particular, species affect one another only through the metabolites they consume and release, so direct interference such as antibiotics, contact-dependent inhibition or signaling is not represented. Uptake of every metabolite saturates with the same half-saturation constant $k$ across all strains and metabolites, and all parameters are fixed for the duration of a simulation, meaning that changes in community behavior arise from shifting abundances and metabolite pools rather than from regulation, time-dependent metabolism or evolution.

##### 1.2 Initial parameter initialization

The initial estimation of parameters in the resource utilization matrix $C_{i\alpha}$, the energy uptake to growth rate conversion $g_i$, and the resource conversion matrix $D^i_{\beta\alpha}$ is made using the exometabolomics measurements, based on a set of simplified ordinary differential equations:

$$\frac{dN_i}{dt} = g_i N_i \sum_\alpha w_\alpha C_{i\alpha} R_\alpha \quad \text{[S3]}$$

$$\frac{dR_\alpha}{dt} = -\sum_i C_{i\alpha} N_i R_\alpha \quad \text{[S4]}$$

With the assumption of a linear growth from resource usage and the absence of nutrient leakage at the initial phase of growth. Using the time-course measurements of exometabolites sampled during the monoculture growth of strain $i$, the resource uptake parameter $C_{i\alpha}$ of resource $\alpha$ can be derived from

$$C_{i\alpha} = max[-\frac{1}{N_{i,t}} \cdot \frac{2(R_{\alpha,t}-R_{\alpha,t-1})}{(R_{\alpha,t}+R_{\alpha,t-1})\cdot dt}] \quad for\ t \in \{T_1, T_2, T_3, T_4\} \qquad [S5]$$

After deriving $C_{i\alpha}$ for each resource $\alpha$, the growth rate conversion $g_i$ is fitted from

$$g_{i,t} = \frac{2(N_{i,t}-N_{i,t-1})}{(N_{i,t}+N_{i,t-1})dt} \cdot \frac{1}{\sum_\alpha w_\alpha C_{i\alpha} R_\alpha} \quad for\ t \in \{T_1, T_2, T_3, T_4\} \qquad [S6]$$

And is shown in Supplementary Fig 5. Additional definitions and units of model parameters can be found in Supplementary Table 3.

The resource conversion matrix $D^i_{\beta\alpha}$ for each strain $i$ describes the secretion of resource $\alpha$ from resource $\beta$ after consumption. At each timepoint that exometabolites were sampled, the secretion of resource $\alpha$ is detected, and the contribution of each resource $\beta$ to the secretion of resource $\alpha$ is normalized based on the consumption level of each resource $\beta$ at that timestep. The final $D^i_{\beta\alpha}$ matrix is averaged at all timesteps and normalized to ensure conservation of mass.

##### 1.3 Simulated annealing algorithm for parameter improvement.

The goal of the algorithm is to find the combination of parameter variables that minimize the score *S* in Equation [3]. Each iteration of the Monte Carlo algorithm generates a new set of the parameters $\{C'_{i\alpha}, D^{i\prime}_{\beta\alpha}, g'_i, l'_{i\alpha}\}$

$$C'_{i\alpha} = C_{i\alpha} + \frac{1}{0.1\cdot C_{i\alpha}\cdot\sqrt{2\pi}} e^{-\frac{1}{2}(\frac{x}{0.1\cdot C_{i\alpha}})^2} \qquad [S7]$$

$$D^{i\prime}_{\beta\alpha} = D^i_{\beta\alpha} + \frac{1}{0.1\cdot D^i_{\beta\alpha}\cdot\sqrt{2\pi}} e^{-\frac{1}{2}(\frac{x}{0.1\cdot D^i_{\beta\alpha}})^2} \qquad [S8]$$

$$g'_i = g_i + \frac{1}{0.1\cdot g_i\cdot\sqrt{2\pi}} e^{-\frac{1}{2}(\frac{x}{0.1\cdot g_i})^2} \qquad [S9]$$

$$l'_{i\alpha} = l_{i\alpha} + 0.01 \cdot \frac{1}{\sqrt{2\pi}} e^{-\frac{1}{2}x^2} \qquad [S10]$$

Based on the current parameters $\{C_{i\alpha}, D^i_{\beta\alpha}, g_i, l_{i\alpha}\}$. For instance, a randomly generated perturbation value, sampled from a Gaussian distribution with 0 mean and $0.1 \cdot C_{i\alpha}$ standard deviation, is added to the current parameter $C_{i\alpha}$ to make the new candidate $C'_{i\alpha}$. Candidate parameters $D^{i\prime}_{\beta\alpha}$ and $g'_i$ are similarly made, while $l'_{i\alpha}$ is generated by adding a randomly drawn value from a standard normal distribution with a factor of 0.01 to the current $l_{i\alpha}$. After simulating the monoculture growth of the given strain, the parameter set score for the new assignment $S'$ is calculated based on equation [3], where $o_1$ and $o_2$ are set to be 10 and 1. For strains *Methylobacterium*, *Marmoricola* and *Paenibacillus*, their respective growth curves measured every 20 minutes on a plate reader spanning 48 hours of culturing period were used to calculate the population abundance difference in simulation and actual measurements. The candidate parameter set is accepted with a probability based on the score change and the temperature:

$$P = e^{-(\frac{S'-S}{T})} \qquad [S11]$$

If the candidate set improved S, then P > 1 and the new candidate set is always accepted. If the candidate set did not improve S, a random number R between 0 and 1 is generated and is compared to the probability P. If R < P, the candidate is accepted and if R > P, the candidate is rejected. The probability of accepting the candidate decreases linearly:

$$T_i = \frac{T_{max}}{i+1} \qquad [S12]$$

where $i$ is the number of iteration, $T_i$ is the temperature factor of the current iteration and $T_{max}$ is the starting temperature. The convergent solutions for all strains in 10 independent runs when $i$ is 10,000 and $T_{max}$ = 100 are then used in the fitted simulations in Fig. 2. Supplementary Fig. 6b-d shows how much these metabolite consumption parameters changed through the simulated annealing step.

### 2 16S data normalizations

#### 2.1 Rationale and scope

Relative abundances we report in polyculture experiments were inferred from 16S rRNA amplicon sequencing. 16S rRNA can be skewed from ground truth cell-count relative abundances due to a wide range of factors that vary between different bacterial strains and experimental protocols including but not limited to 16S rRNA copy number, cell lysis efficiency, amplification efficiency of 16S gene sequence, primer binding efficiency, and number of PCR cycles. In the analysis reported in the main text, sequencing read fractions are used without correction. This supplementary section in addition to Supplementary Fig. 9 and 10 describes (i) the possible normalizations we could and did implement from our data and (ii) the robustness of the main model-experiment comparisons to alternative model choices, specifically in the context of the whole community assembly experiment (Fig. 4a-e).

#### 2.2 16S copy number

16S rRNA gene copy numbers for each isolate were retrieved from the supplementary data of Novak et al. 2025, which reports genome information for the members of the model switchgrass-rhizosphere SynCom. In that study, whole genomes were sequenced at the Joint Genome Institute on either the PacBio Sequel II or the Illumina NovaSeq S4 platform. Assemblies were annotated through the Integrated Microbial Genomes and Microbiomes (IMG/M) pipeline; 16S rRNA gene copy numbers were taken from these annotations. Annotated genomes are accessible via Hugging Face (https://doi.org/10.57967/hf/5885).

Across the isolates used in this study, 16S rRNA gene copy number ranged from 1 to 13 per genome. Copy numbers for each strain are given in Supplementary Table 5.

In Supplementary Fig. 10d when 16S copy number is normalized for the correction procedure is as follows:

$$\text{Normalized Abundance} = \frac{\text{Raw 16S Relative Abundance}}{\text{16S rRNA gene copy number}} \quad [S13]$$

##### 2.3 Strain-specific OD-to-CFU conversion factors

Communities were inoculated by mixing strains at equal optical density (OD). Because the relationship between OD and cell number is strain-specific, equal-OD inoculation does not correspond to equal cell number. We therefore measured strain-specific OD-to-CFU conversion factors for our initial inoculation OD of 0.1 to allow initial cell fractions to be estimated independently of sequencing.

Strains were streaked from frozen stock onto 1X NLDM 1.5% agar plates and incubated for 48hr at 30°C. Individual colonies were used to inoculation starter cultures in 3mL of 1X NLDM in a 15 mL culture tube (Sigma, CLS352057). Starter cultures were prepared in biological triplicate cultures and included negative control tubes (no cells). Cultures were grown for 48hr in a 30° C incubator with shaking (Fisher, MaxQ Incubated Shaker).

On Day 0, individual colonies from the plates were used to inoculation starter cultures in 3mL of 1X NLDM in a 15 mL culture tube (Sigma, CLS352057). Starter cultures were prepared in biological triplicate cultures and included negative control tubes (no cells). Cultures were grown for 48hr in a 30° C incubator with shaking.

On Day 1, 1x NLDM + 1.5% agar plates were prepared to run the plating experiment for strains in biological duplicate, with technical plating triplicates, at three serial dilution concentrations (i.e. 18 plates per strain). It is recommended to label plates with strain information and dilution information prior to beginning the plating process on Day 2.

On Day 2, the starter cultures were serially diluted (factor of 10) in a 96 well plate to determine what target dilution would give an OD600nm of ~0.1-0.2. A plating stock with a target OD600nm of 0.1 was then prepared for each strain based on the OD600 findings. Plating stocks were then serially diluted in a 96 well plate by using a multichannel pipette to serially transfer 20uL of plating stock into 180uL of fresh 1X NLSM media, generating a dilution series from 10^-1 to 10^-7. For plating, 4-5 sterile glass beads were added to each labeled plate, 50uL of the dilution ($10^{-4}$ and $10^{-5}$ and $10^{-6}$) was pipetted onto each plate, and then plates were covered and gently shaken in a circular pattern to distribute the inoculum evenly across the plate. Spent beads were poured into a waste beaker for reuse. This protocol was used for each strain (18 plates per strain). Plates were incubated upside down at 30°C and observed daily for the next 7 days.

When individual colonies on plates grew large enough to be distinguishable by eye, images of the plates were captured with a Nikon digital SLR camera. Images were processed using Fiji/ImageJ for counting, and only those with a colony density between 30-300 CFU were considered viable for analysis. OD-to-CFU (in CFU/mL at OD 0.1) was calculated according to Equation S14, where C is the sum number of counts across technical triplicates, V is the volume plated across technical triplicates, and D is the serial dilution factor used for plating. For each count for each dilution, a pooled standard deviation was quantified across biological replicates (where s is the standard deviation for each biological replicate and k is the total number of plates (Equation S15) included in the average).

$$\text{OD-to-CFU } \left(\frac{CFU}{mL} \text{ at OD } 0.1\right) = \frac{\sum C}{\sum V*D} \pm \frac{\sqrt{\sum C}}{\sum V*D} \quad \text{[S14]}$$

$$SD_{pooled} = \sqrt{\frac{s_1^2 + s_2^2 + \cdots s_k^2}{k}} \quad \text{[S15]}$$

Mean and standard deviations of this calculated OD-to-CFU across dilutions and replicates can be found reported for each strain in Supplementary Table 5. For a small number of strains, serial dilutions did not yield plates within the acceptable counting range (30–300 CFU per plate), so no strain-specific conversion factor could be determined. For these strains the default conversion of $1 \times 10^9$ CFU $mL^{-1}$ at OD 0.1 was used for these additional analyses instead.

These factors were used to adjust the initial conditions in our simulations (Supplementary Fig. 10e) such that instead of equal starting abundances across strains times a default $1 \times 10^9$ CFU $mL^{-1}$ conversion, the conversion used was the strain specific ones reported in Supplementary Table 5.

##### 2.4 Explanatory power of 16S copy number and OD-to-CFU conversion for time-zero composition

For one replicate community in experiment B (Fig. 4a), we sequenced the community immediately after inoculation. Because strains were combined at equal OD, each of the $N$ strains should account for a relative abundance of $1/N$ at this point, so any departure from that should reflect measurement bias rather than any biological process.

We regressed this observed time-zero relative abundance against 16S copy number, and OD-to-CFU, and the product of the two, using a simple linear regression across the 15 strains in the community (Supplementary Fig. 10a-c). None of the three predictors explained a substantial fraction of the variation (16S copy #: $R^2 = 0.01$, $p = 0.734$; CFU/mL: $R^2 = 0.00$, $p = 0.862$; both: $R^2 = 0.02$, $p = 0.646$), indicating that additional unmeasured factors also contribute.

We therefore used the time-zero composition itself as a measure of the combined bias acting on each strain, defining a deviation factor:

$$\text{Deviation factor(i)} = \frac{p_i^0}{1/N} \quad \text{[S16]}$$

where $p_i^0$ is the time-zero raw 16S-derived relative abundance of strain $i$ normalized to sum to one across strains. Values above and below 1 indicate strains respectively over- and under-represented relative to what was inoculated. Strains not detected at time zero were assigned a deviation factor of 1. Deviation factors are reported in Supplementary Table 5.

##### 2.5 Alternative normalizations applied to experimental relative abundances

To test whether the main experiment-model comparisons depend on normalization choice, we recomputed community compositions under two alternative schemes and repeated the analyses of the main text.

First we normalized the raw 16S data by 16S copy number by dividing the relative abundances by that strains specific copy number as in [S13]. Second, we normalized the composition of the raw16S output by the deviation factor calculated from the time-zero 16S sequencing when strains were mixed at equal OD as in [S16]. Each of these two normalizations were performed across both whole community experiments and all passages (Supplementary Fig. 10d).

To evaluate quantitatively how these normalizations affect experiment-model comparisons, we performed the same correlation analysis as the main manuscript (Fig. 4e) and compared the Spearman rank correlation coefficient across the various choices of normalizations across passages (Supplementary Fig. 9a) and at the final sampling timepoint at the end of the experiments (Supplementary Fig. 9b). Normalization had only marginal effects on the correlations and preserved the key contrasts (Experiment A versus B; model with versus without cross-feeding), so our conclusions are unaffected and we show raw 16S data in Fig. 4.

##### 2.6 Alternative initial conditions in the consumer–resource model

In the CRM implementation reported in the main text, all strains are initialized at equal abundance, based on the equi-OD initialization in the experiment. One could alternatively choose the 16S-derived relative abundance (or other normalized versions of it) as initial condition, and compare results across these different initializations. Specifically, to assess the sensitivity of CRM predictions to strain initial conditions (in addition to experimental data normalizations), we repeated the whole-community simulations under three alternative initializations: (1) initial abundances set to the measured time-zero 16S composition (2) initial abundances scaled by strain-specific OD-to-CFU conversion factors and (3) both adjustments combined. Across all initializations, the no cross-feeding scenario remained clearly distinct from the cross-feeding scenarios (Supplementary Fig. 10e), and correlations with experimental abundances remained close to those reported in the main text (Supplementary Fig. 9). A more thorough exploration of the sensitivity of the simulated results to initial conditions is also performed and shown in Supplementary Fig. 11, and described in detail below.

#### 3 Initial community abundance perturbations

To understand the effects of initial community abundances on our model, we varied these quantities in across 100 different simulations, which are shown in Supplementary Fig. 11a.

To vary initial community abundances, time zero 16s relative abundances by strains were shuffled and applied as the initial community abundances as described above. Whole-community simulations were done as described in the main methods.

The distributions of final abundances of each strain for varied initial community compositions are shown in Supplementary Fig. 11b, which are plotted in tandem with final community abundances as simulated based on different normalization techniques. As these all land within the observed distributions, we can conclude that initial strain conditions have minimal effect on simulated community outcome.

#### 4 Computing p-value for pairwise performance

##### 4.1 Rationale and scope

In our pairwise strain prediction validation, we compare to data with two experimental replicates, which limits the statistical tests we can perform. Besides showing $R^2$ and RMSE values calculated using the sklearn.metrics package, we also performed a label-shuffling method allowing us to better quantify the performance of our model compared to one agnostic to strain-level information, highlighted in Supplementary Fig 8b.

##### 4.2 Dataset permutations

16s results were represented as a 15 x 15 matrix with paired ratios – each column represents one strain relative abundance in pairwise co-culture with the strain on each row. For each permutation, the shared labels of rows and columns were shuffled such that experimental information was maintained but strain information was randomized. The $R^2$ for the resulting labels was calculated compared to the unshuffled simulation results, and p value was calculated by dividing the number of permutations with $R^2$ equal to or greater than the observed value divided by the total number of permutations done (9999).

#### 5 Carbon usage in exometabolomics data

In previous methods, handling of metabolomics data is described. To calculate total carbon usage of individual strains in monoculture, normalized values were multiplied by concentration of metabolites in the media in µM and then by carbon counts derived from molecular formulas. Secretion was calculated by adding up carbon added to the media compared to initial media concentration, and consumption was calculated by adding up carbon consumed compared to initial condition. Net flux is defined as change in µM from the initial media condition. All results are shown in Supplementary Fig. 3.

### Supplementary Tables

**Supplementary Table 1.** Strains in 15-member synthetic community, with strain names and GenBank Assembly IDs specified and reported plants known to be host partners. Individual strains can be found on DSMZ or as a collection (DSM 200000SY, inquire at). Some species have alternative names, listed in parentheses. Table modified from Coker (2022).

| Genus | Strain | GenBank Assembly | Associated plants |
|---|---|---|---|
| *Lysobacter* | OAE881 | GCA_040546665.1 | Nicotiana tabacum, tomato, pepper |
| *Burkholderia (Paraburkholderia)* | OAS925 | GCA_040543975.1 | Zea mays, Betula, Equisetum, Quercus, Senecio vulgaris, Triticum aestivum, Zantedeschia, Coffea, Saccharum officinarum, Lolium multiflorum, citrus, wheat |
| *Variovorax* | OAS795 | GCA_040546685.1 | Citrus, maize, tomato, wheat |
| *Chitinophaga* | OAE865 | GCA_040546735.1 | Tomato, Oryza sativa, Cymbidium goeringii, ginseng |
| *Niastella* | OAS944 | GCA_040546725.1 | Hibiscus syriacus, persimmon tree, Populus euphratica, Korean ginseng |
| *Mucilaginibacter* | OAE612 | GCA_041806395.1 | Gossypium hirsutum, Angelica sinensis, ginseng, Dokdo Island (S Korea) |
| *Rhizobium* | OAE497 | GCA_040546565.1 | Citrus, Oryza sativa, Dioscorea alata, Dioscorea esculenta |
| *Bosea* | OAE506 | GCA_040546595.1 | Zea mays, Cyperus rotundus |
| *Methylobacterium* | OAE516 | GCA_041806315.1 | Eucalyptus spp., Oryza sativa cv. Dongjin and Lycopersicon esculentum. cv. Mairoku, maize |
| *Arthrobacter* | OAP107 | GCA_040546765.1 | Ginkgo biloba, Quercus ilex, Triticum aestivum(wheat) |
| *Rhodococcus* | OAS809 | GCA_040546695.1 | Zea mays, Oryza sativa |
| *Brevibacillus* | OAP136 | GCA_040544025.1 | Zea mays, Lolium perenne, Pinellia ternata, Nicotiana tabacum, Gossypium hirsutum |
| *Paenibacillus* | OAE614 | GCA_040546645.1 | Solanum lycopersicum, Oryza sativa, wheat |
| *Bacillus (Gottfriedia)* | OAE603 | GCA_040543965.1 | Zea mays, Triticum aestivum (wheat), Lolium perenne, Nicotiana tabacum |
| *Marmoricola* | OAE513 | GCA_040546585.1 | Kandelia candel (mangrove), Camellia sinensis, Papaver somniferum, Brassica chinensis, wheat |

**Supplementary Table 2.** List of 64 metabolites in the Northern Lab Defined Medium (NLDM) used in Consumer-Resource Model from Targeted Exometabolomics profiling

| Name | Class | Formula | Concentration (uM) |
|---|---|---|---|
| Sucrose | sugar, dihexose | $C_{12}H_{22}O_{11}$ | 875 |
| Glucose | sugar, hexose | $C_6H_{12}O_6$ | 875 |
| Trehalose | sugar, dihexose | $C_{12}H_{22}O_{11}$ | 875 |
| Myo-inositol | sugar, hexose alcohol | $C_6H_{12}O_6$ | 875 |
| Xylose | sugar, pentose | $C_5H_{10}O_5$ | 875 |
| Mannitol | sugar, hexose alcohol | $C_6H_{14}O_6$ | 875 |
| n-Acetyl glucosamine | sugar, amino sugar | $C_8H_{15}NO_6$ | 875 |

| | | | |
|---|---|---|---|
| Sodium-L-Lactate | organic acid | $C_3H_5O_3Na$ | 525 |
| Malic acid | organic acid | $C_4H_6O_5$ | 525 |
| Citric acid | organic acid | $C_6H_8O_7$ | 525 |
| Succinic acid | organic acid | $C_4H_6O_4$ | 525 |
| Pyruvic acid | organic acid | $C_3H_4O_3$ | 525 |
| Alpha-Ketoglutaric acid | organic acid | $C_5H_6O_5$ | 525 |
| Citrulline | amino acid | $C_6H_{13}N_3O_3$ | 525 |
| Alanine | amino acid | $C_3H_7NO_2$ | 175 |
| Arginine | amino acid | $C_6H_{14}N_4O_2$ | 175 |
| Asparagine | amino acid | $C_4H_8N_2O_3$ | 175 |
| Aspartic acid | amino acid | $C_4H_7NO_4$ | 175 |
| Cystine/Cysteine | amino acid | $C_3H_7NO_2S$ | 175 |
| Glutamic acid | amino acid | $C_5H_9NO_4$ | 175 |
| Glutamine | amino acid | $C_5H_{10}N_2O_3$ | 175 |
| Glycine | amino acid | $C_2H_5NO_2$ | 175 |
| Histidine | amino acid | $C_6H_9N_3O_2$ | 175 |
| Isoleucine | amino acid | $C_6H_{13}NO_2$ | 175 |
| Leucine | amino acid | $C_6H_{13}NO_2$ | 175 |
| Lysine | amino acid | $C_6H_{14}N_2O_2$ | 175 |
| Methionine | amino acid | $C_5H_{11}NO_2S$ | 175 |
| Phenylalanine | amino acid | $C_9H_{11}NO_2$ | 175 |
| Proline | amino acid | $C_5H_9NO_2$ | 175 |
| Serine | amino acid | $C_3H_7NO_3$ | 175 |
| Threonine | amino acid | $C_4H_9NO_3$ | 175 |
| Tryptophan | amino acid | $C_{11}H_{12}N_2O_2$ | 175 |
| Tyrosine | amino acid | $C_9H_{11}NO_3$ | 175 |
| Valine | amino acid | $C_5H_{11}NO_2$ | 175 |
| 5-methyluridine | nucleoside | $C_{10}H_{14}N_2O_6$ | 17.5 |
| Adenine | nucleoside | $C_5H_5N_5$ | 17.5 |
| Adenosine | nucleoside | $C_{10}H_{13}N_5O_4$ | 17.5 |
| Cytidine | nucleoside | $C_9H_{13}N_3O_5$ | 17.5 |
| Cytosine | nucleoside | $C_4H_5N_3O$ | 17.5 |
| Guanine | nucleoside | $C_5H_5N_5O$ | 17.5 |
| Guanosine | nucleoside | $C_{10}H_{13}N_5O_5$ | 17.5 |
| Hypoxanthine | nucleoside | $C_5H_4N_4O$ | 17.5 |
| Inosine | nucleoside | $C_{10}H_{12}N_4O_5$ | 17.5 |
| Thymine | nucleoside | $C_5H_6N_2O_2$ | 17.5 |
| Uracil | nucleoside | $C_4H_4N_2O_2$ | 17.5 |
| Uridine | nucleoside | $C_9H_{12}N_2O_6$ | 17.5 |
| Xanthine | nucleoside | $C_5H_4N_4O_2$ | 17.5 |
| Xanthosine | nucleoside | $C_{10}H_{12}N_4O_6$ | 17.5 |

| | | | |
|---|---|---|---|
| gamma-Aminobutyric acid | nucleoside | $C_4H_9NO_2$ | 17.5 |
| Ectoine | osmolyte | $C_6H_{10}N_2O_2$ | 17.5 |
| Trimethylglycine | amino acid derivative | $C_5H_{11}NO_2$ | 17.5 |
| N-Acetyl-glutamic acid | amino acid derivative | $C_7H_{11}NO_5$ | 17.5 |
| Nicotinamide | vitamin | $C_6H_6N_2O$ | 17.5 |
| Ornithine HCl | amino acid derivative | $C_5H_{12}N_2O_2HCl$ | 17.5 |
| Shikimic acid | other | $C_7H_{10}O_5$ | 17.5 |
| Taurine | other | $C_2H_7NO_3S$ | 17.5 |
| Trigonelline HCl | alkaloid | $C_7H_7NO_2HCl$ | 17.5 |
| Carnitine HCl | other | $C_7H_{16}NO_3HCl$ | 17.5 |
| choline chloride | osmolyte | $C_5H_{14}NOCl$ | 17.5 |
| n-Acetyl-glutamine | amino acid derivative | $C_7H_{12}N_2O_4$ | 17.5 |
| n-Acetyl-lysine | amino acid derivative | $C_8H_{16}N_2O_3$ | 17.5 |
| n-Acetyl-muramic acid | other | $C_{11}H_{19}NO_8$ | 17.5 |
| sn-glycero-3-phosphocholine | phospholipid | $C_8H_{20}NO_6P$ | 17.5 |

**Supplementary Table 3**. Definitions and units of consumer resource model dynamical variables and mechanistic parameters.

| **Variable/parameter** | **Description** | **Units** | **Quantity** |
|---|---|---|---|
| $N_i$ | Abundance of organism i | CFU/mL | Variable |
| $R_\beta$ | Abundance of resource β | g/mL | Variable |
| $C_{i\alpha}$ | Uptake rate per unit concentration of resource α | mL/hr | Variable |
| $D_{\alpha\beta}^{i}$ | Proportion of resource β converted to resource α by organism i | Unitless | Variable |
| $l_{i\alpha}$ | Leakage fraction of resource α in organism i | Unitless | Variable |
| $g_i$ | Conversion factor from energy uptake to growth rate for organism i | 1/energy | Variable |
| $w_\alpha$ | Energy content of resource α | energy/g | $1 \times 10^{12}$ |
| $k_{i\alpha}$ | Half velocity constant for resource uptake | g/mL | 0.04 |

**Supplementary Table 4**. Selected 3-strain subcommunities for experimental validation

| Community Index | Top 10 selected community | Community Index | Bottom 10 selected community |
|---|---|---|---|
| 1 | *Burkholderia, Mucilaginibacter, Bacillus* | 11 | *Variovorax, Chitinophaga, Paenibacillus* |
| 2 | *Variovorax, Brevibacillus, Paenibacillus* | 12 | *Niastella, Arthobacter, Paenibacillus* |
| 3 | *Burkholderia, Rhizobium, Bacillus* | 13 | *Burkholderia, Bosea, Paenibacillus* |
| 4 | *Mucilaginibacter, Rhizobium, Bacillus* | 14 | *Bosea, Paenibacillus, Bacillus* |
| 5 | *Burkholderia, Mucilaginibacter, Rhizobium* | 15 | *Burkholderia, Variovorax, Bosea* |

| | | | |
|---|---|---|---|
| 6 | *Lysobacter, Niastella, Rhodococcus* | 16 | *Variovorax, Bosea, Arthobacter* |
| 7 | *Burkholderia, Chitinophaga, Rhizobium* | 17 | *Variovorax, Mucilaginibacter, Bosea* |
| 8 | *Lysobacter, Chitinophaga, Niastella* | 18 | *Bosea, Arthobacter, Brevibacillus* |
| 9 | *Chitinophaga, Niastella, Rhodococcus* | 19 | *Variovorax, Chitinophaga, Bosea* |
| 10 | *Chitinophaga, Mucilaginibacter, Rhodococcus* | 20 | *Arthobacter, Brevibacillus, Paenibacillus* |

**Supplementary Table 5**. Strain-specific data for 16S sequencing correction analysis

| **Strain** | **16S rRNA Copy Number** | **Deviation Factor (T0 sequencing)** | **OD-to-CFU mean (CFU mL$^{-1}$)** | **OD-to-CFU std (CFU mL$^{-1}$)** |
|---|---|---|---|---|
| *Lysobacter* | 2 | 0.30 | $1.0 \times 10^9$ | $5.0 \times 10^8$ |
| *Burkholderia* | 6 | 1.42 | $1.1 \times 10^8$ | $1.3 \times 10^7$ |
| *Variovorax* | 1 | 0.59 | $2.4 \times 10^8$ | $2.8 \times 10^8$ |
| *Chitinophaga* | 6 | 4.65 | $6.3 \times 10^8$ | $1.9 \times 10^8$ |
| *Niastella* | 4 | 2.01 | $7.4 \times 10^7$ | $1.7 \times 10^7$ |
| *Mucilaginibacter* | 3 | 2.01 | $4.6 \times 10^8$ | $4.5 \times 10^7$ |
| *Rhizobium* | 4 | 1.27 | $1.6 \times 10^7$ | $2.0 \times 10^6$ |
| *Bosea* | 2 | 0.054 | $1.0 \text{ x } 10^9$ | Default conversion |
| *Methylobacterium* | 4 | 0.024 | $9.4 \times 10^6$ | $1.1 \times 10^6$ |
| *Arthrobacter* | 6 | 0.16 | $5.1 \times 10^7$ | $2.0 \times 10^7$ |
| *Rhodococcus* | 5 | 0.40 | $1.4 \times 10^8$ | $1.4 \times 10^8$ |
| *Brevibacillus* | 13 | 1.0 | $1.0 \text{ x } 10^9$ | Default conversion |
| *Paenibacillus* | 9 | 1.0 | $1.3 \times 10^6$ | $1.3 \times 10^5$ |
| *Bacillus* | 11 | 2.11 | $1.0 \text{ x } 10^9$ | Default conversion |
| *Marmoricola* | 1 | 1.0 | $1.0 \text{ x } 10^9$ | Default conversion |

### Supplementary Figures

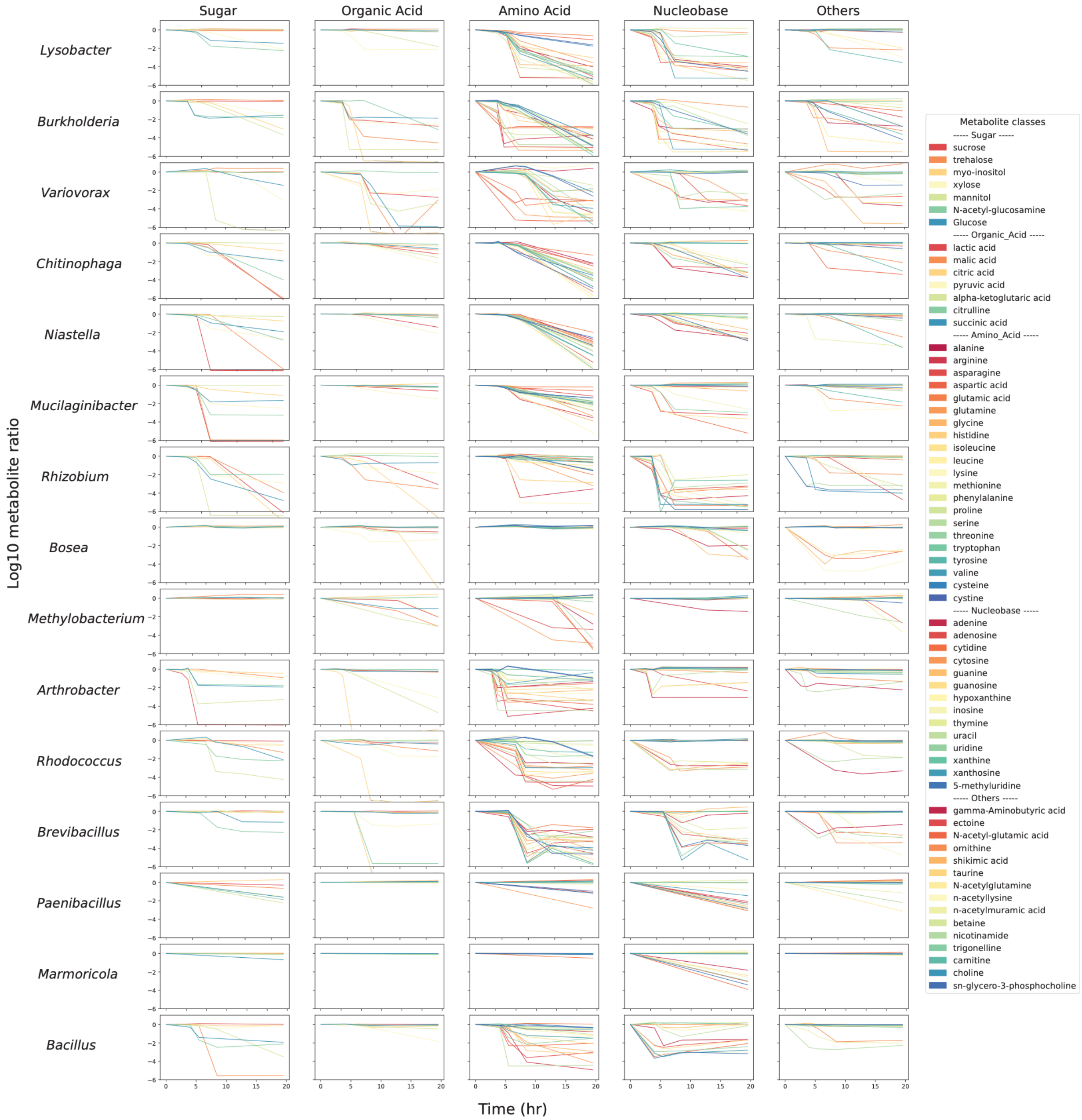


**Supplementary Figure 1. Metabolite concentrations over time in monoculture experiments.** Plotted in the log ratio of the metabolite concentration in monoculture and that metabolite concentration in blank media. A value of 0 corresponds to no net consumption or production of metabolite, positive values show production, and negative values denote consumption.

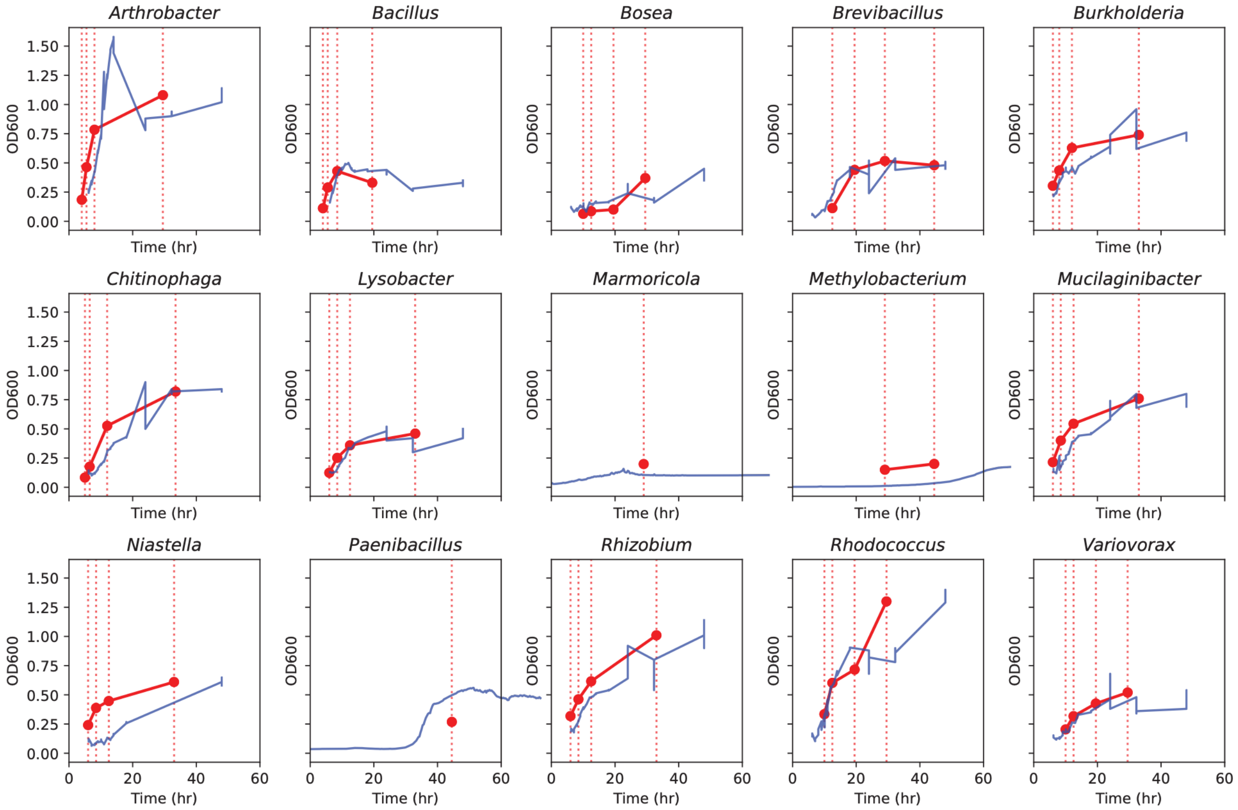


**Supplementary Figure 2. Monoculture measurements of growth over time**. OD growth is shown in blue with the timepoints where exometabolomics measurements were taken (vertical lines). OD measurements were also taken during the 4 sampling points when exometabolomics samples were taken, indicated by red lines. High-density OD measurements (blue) and exometabolomics (red) experiments were done separately.

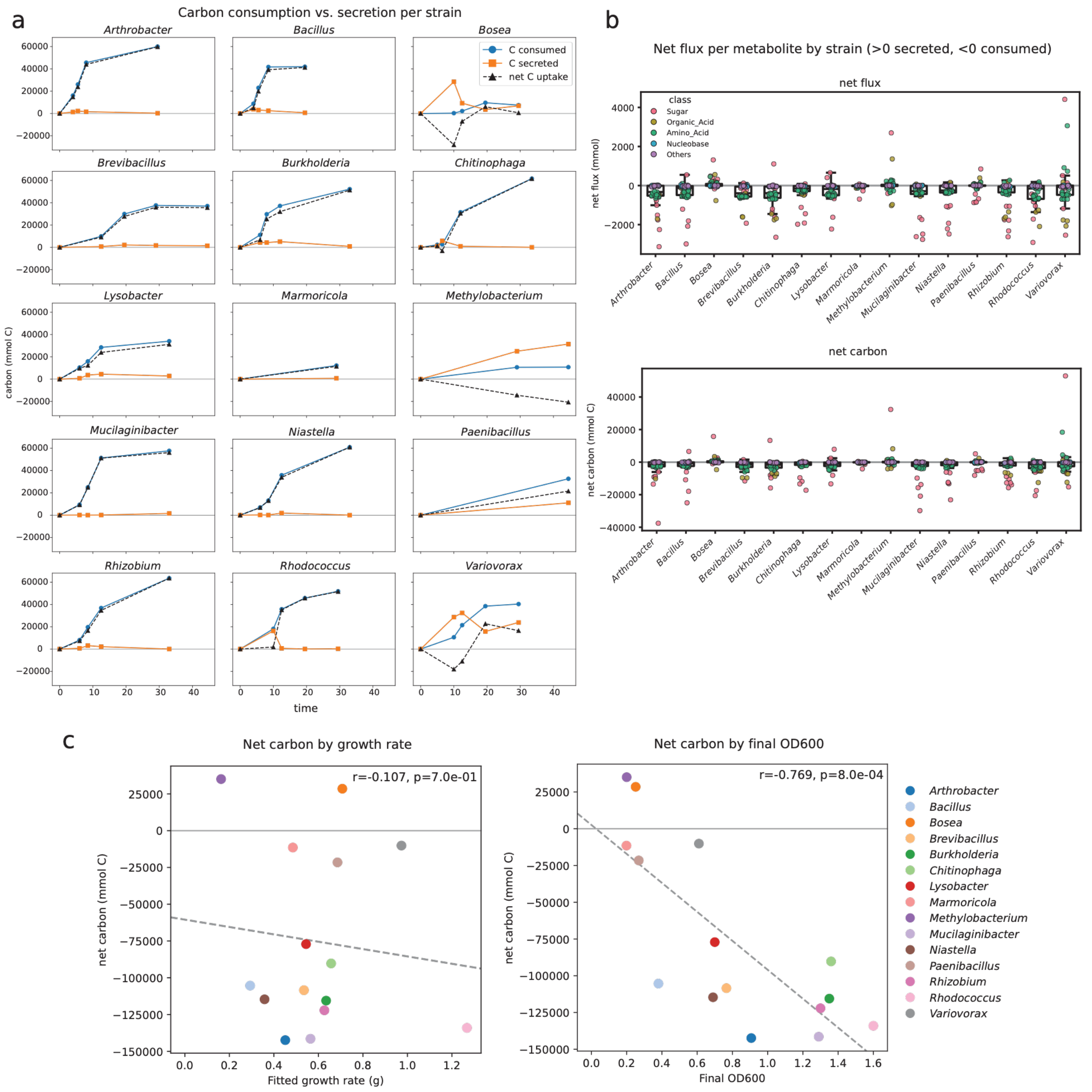


**Supplementary Figure 3. Carbon consumption and secretion varies by strain.** a) Plot of total carbon uptake or secretion by strain. b) Distribution of total carbon and total flux per metabolite plotted by strain. Metabolite class indicated by face color of points consistent with Fig. 2 (pink: sugar, yellow: organic acid, green: amino acid, blue: nucleobase, purple: others). c) Linear fit of growth rate by net carbon uptake (left) and final OD by net carbon uptake (right).

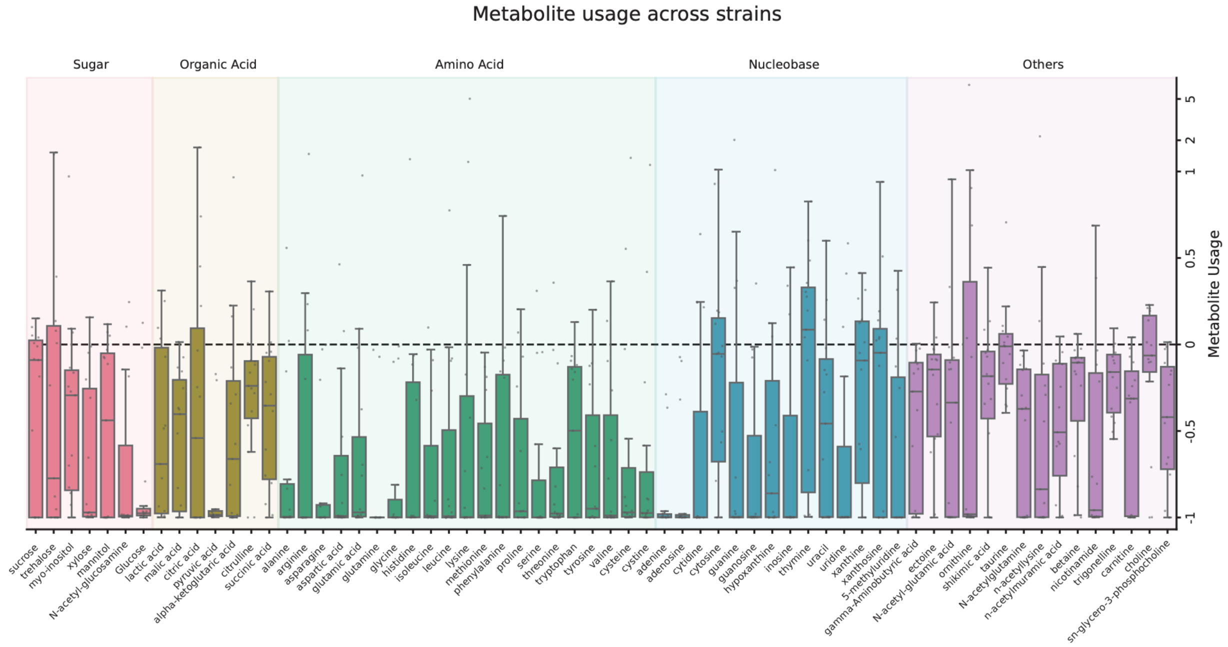


**Supplementary Figure 4. Metabolite usage varies widely across strains for most metabolites.** Distribution of metabolite usage across the 15 monoculture strains at the final timepoint, for each of the 64 metabolites measured (the same data shown as a heatmap in Fig. 2a). Negative values indicate net consumption of a metabolite relative to the medium control, positive values net production. Boxes span the interquartile range with the median marked. Individual strains are overlaid as points (n = 15 per metabolite), so all values are visible, including those beyond the whiskers. Rows are grouped and colored by metabolite class (sugars, organic acids, amino acids, nucleobases, and others), in the same order as Fig. 2a. The x axis is linear between −1 and 1 and logarithmic beyond, so that a small number of strongly accumulating cases (e.g. ornithine, lysine) do not compress the remaining distributions.

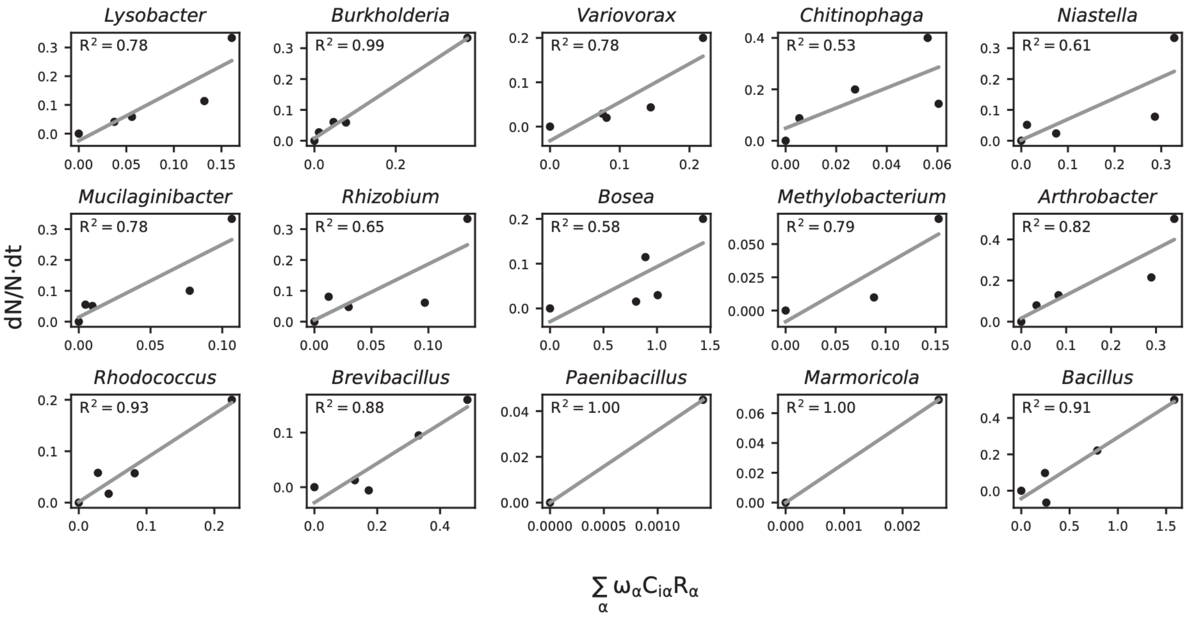


**Supplementary Figure 5. Fitting of the growth parameter $g_i$, shown here as the slope.** For each strain *i*, its $g_i$ parameter is calculated from Equation 2 after inferring the $C_{i\alpha}$ parameter for each metabolite α. $R^2$ for each $g_i$ parameter fitting is shown in individual subplots.

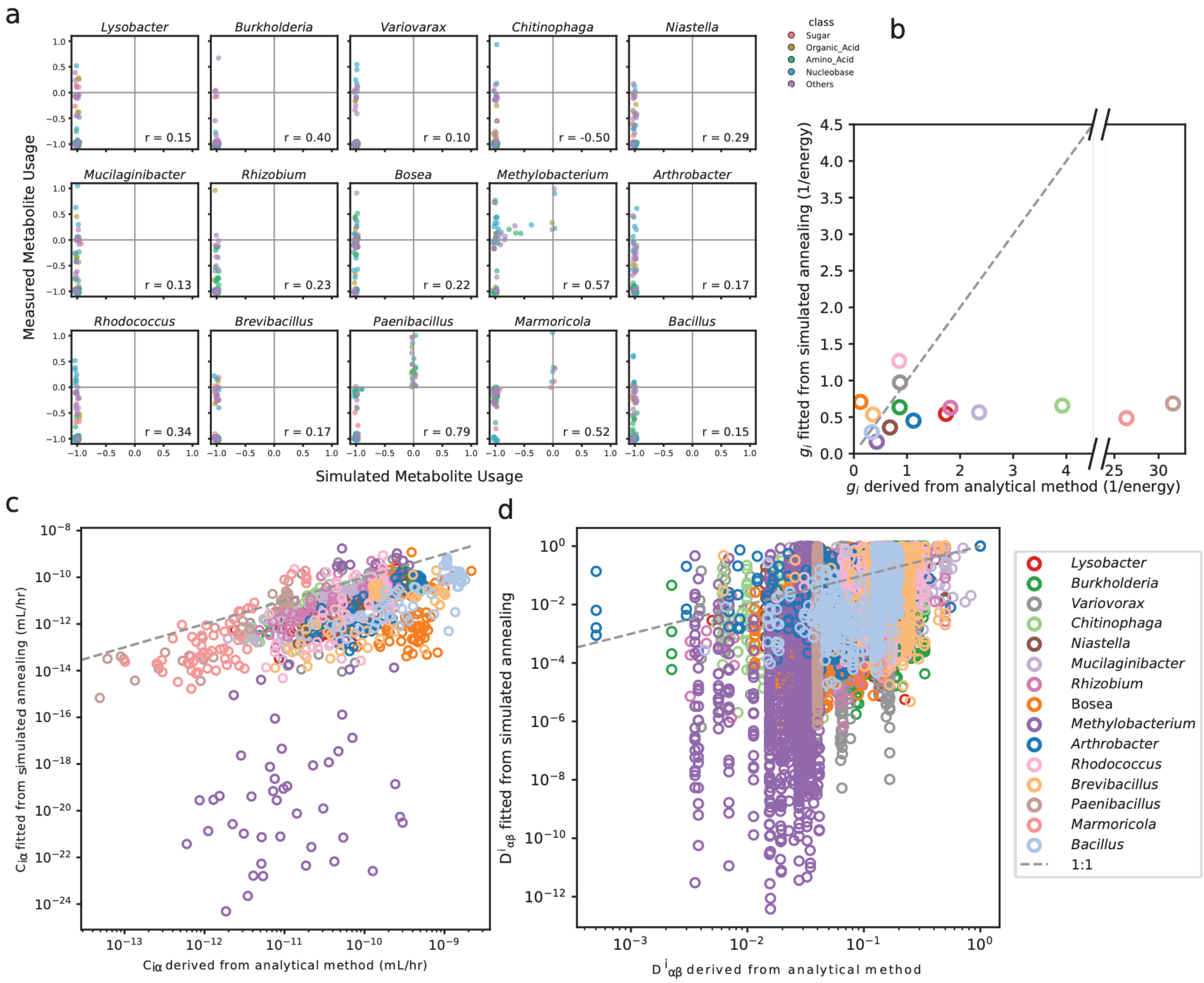


**Supplementary Figure 6. Initial and fitted CRM parameter comparison.** a) Comparison of resource utilization and secretion at the final sampling point determined by exometabolomics measurements and by model prediction using the initial parameters. A positive value denotes resource production and a negative value denotes resource secretion normalized by metabolite initial concentrations. Different classes of molecules are shown in different colors.

b) Comparison of the initial analytically derived growth parameters against the parameters obtained from simulated annealing. Points are colored by species.

c) Comparison of the initial analytically derived metabolite uptake parameters (C matrix) against the parameters obtained from simulated annealing. Points are colored by species.

d) Comparison of the initial analytically derived metabolite transformation parameters (D matrix) against the parameters obtained from simulated annealing. Points are colored by species.

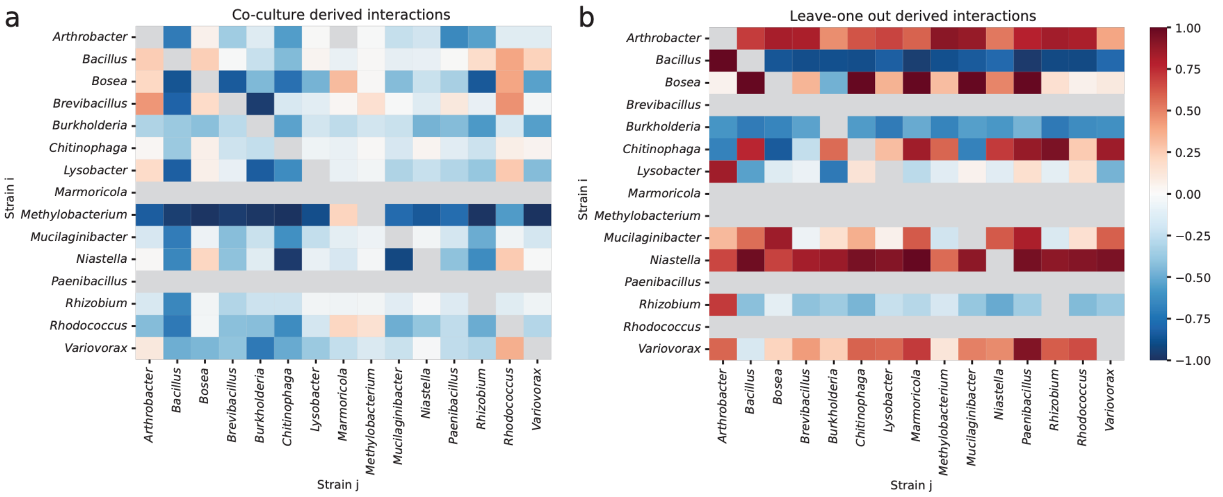


**Supplementary Figure 7. Interaction metrics defined by co-culture versus leave-out experiments.** a) Pairwise strain interactions inferred from co-culture experiments. Conditions were run where the same strain was inoculated twice instead of two different strains for the co-culture, so these interactions are inferred from the final OD measurements and relative abundance 16S data. Fitness of strain i in coculture with strain j ($W_{ij}$) and fitness of strain i without strain j ($W_{ii}$), were then combined into the metric $R_{ij} = W_{ij} – W_{ii} / W_{ij} + W_{ii}$. A positive interaction indicates that strain i grows to a higher population abundance in the presence of j than without. b) Pairwise strain interactions inferred from leave-one-out experiments. In this case, the fitness of strain i when strain j is left out of the community is $W_{i\text{-}j}$ and this is compared to the fitness of i in the whole community $W_{ij*}$ (rescaled with the relative abundance of j removed). A similar metric can then be derived with $R_{ij} = W_{ij*} – W_{i\text{-}j} / W_{ij*} + W_{i\text{-}j}$. The final $R_{ij}$ is positive when the removal of strain j is detrimental to the fitness of strain i. Grey boxes indicate samples with zero/near-zero growth in the whole community context for strain i.

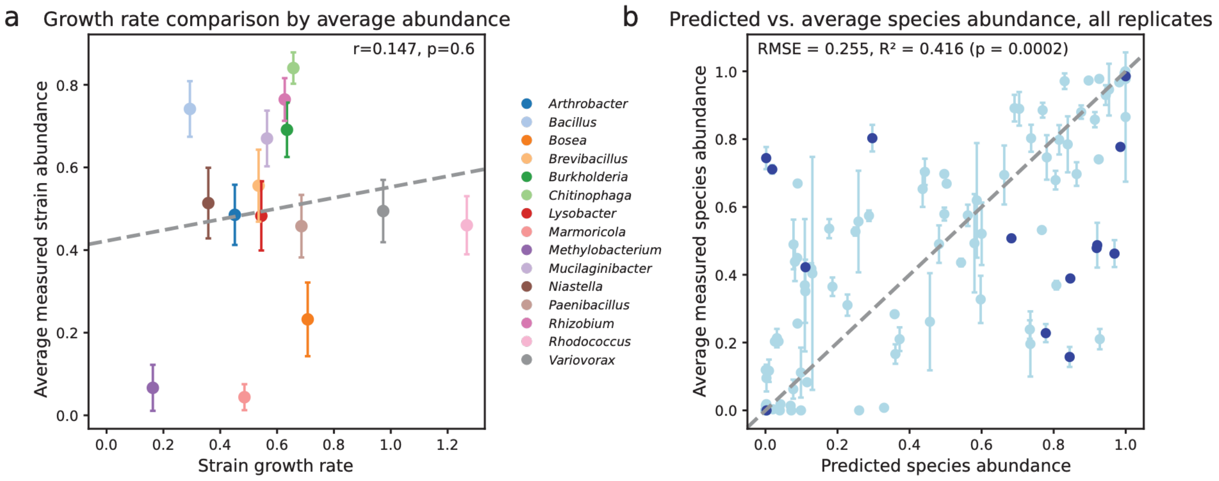

**Supplementary Figure 8. Co-culture strain ratios are predicted by CRM, not by growth rate alone.** a) CRM fitted growth parameters ($g_i$) from monoculture growth are plotted against average co-culture abundances across strains to show that growth is not a sufficient predictor or abundances in co-culture. Line of best fit and pearson correlation results reported. b) Across all co-culture pairs, the ratio of final strain abundances was taken and plotted the simulated results from the fitted CRM and the experimental 16S final strain abundances. RMSE is reported as well as R2. See Supplementary Methods 4 for information about p-values. Pairs involving *Arthrobacter* are colored in dark blue and show higher deviations between experiments and simulations than other strains.

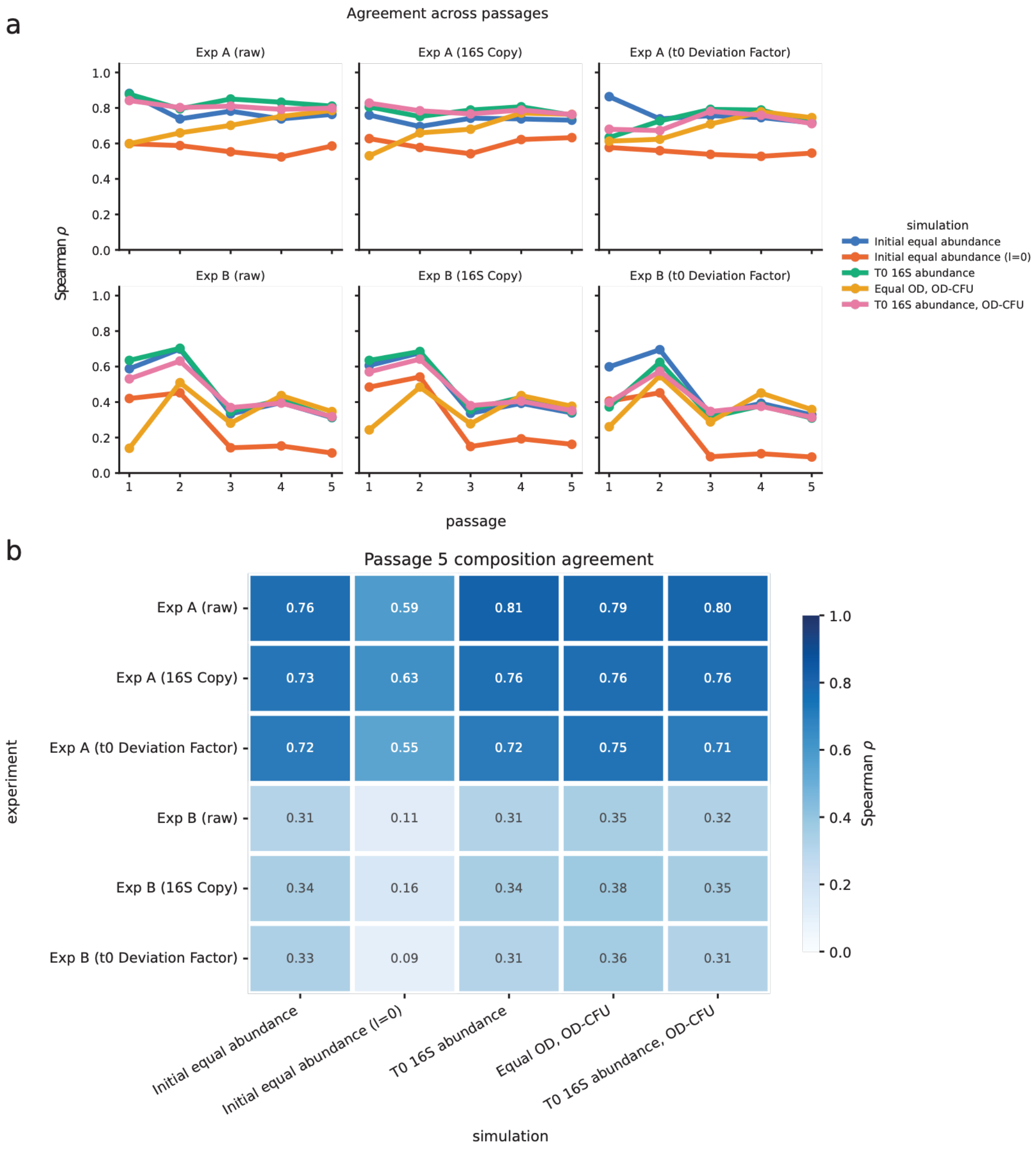


**Supplementary Figure 9. Effects of 16S normalization schemes on whole community correlation results.** a) Spearman rank correlation (ρ) between measured and simulated strain relative abundances at each passage, for each combination of experiment and normalization (panels) and simulation initialization (colors). Agreement is consistently high for Exp A (ρ ≈ 0.6-0.9) and lower and less stable for Exp B (ρ ≈ 0.1-0.7, declining after passage 2). Within each panel the initialization schemes track one another closely, except the l = 0 (no cross-feeding) case, which performs worst throughout. Details on the various normalization schemes can be found in the Supplementary Methods 2. b) Spearman ρ between measured and simulated composition at passage 5 for all experiment × normalization (rows) and initialization (columns) combinations. Values range from 0.09 to 0.81. Agreement is governed primarily by which experiment is considered and whether cross-feeding is represented, rather than by the normalization or initialization applied.

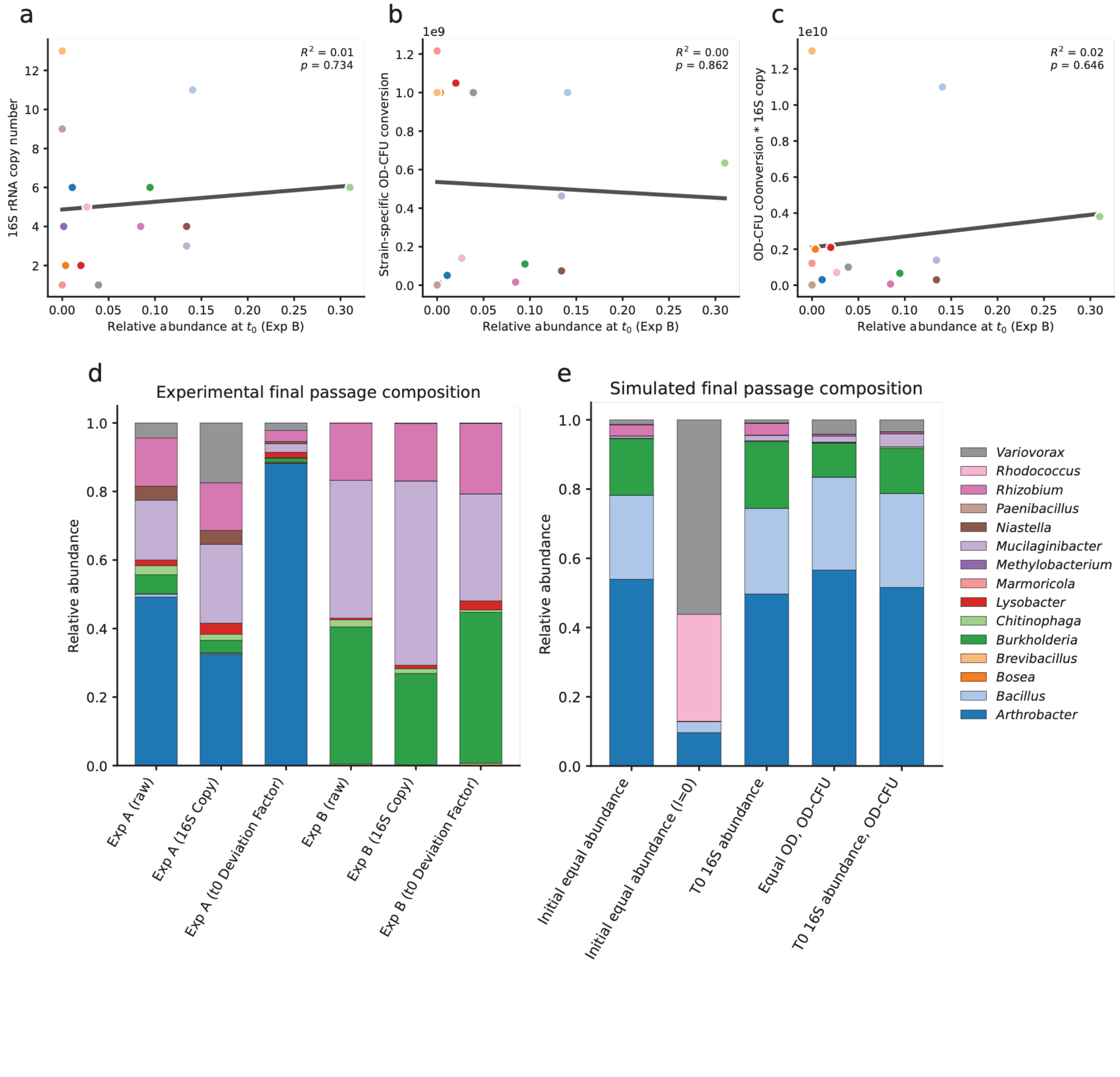


**Supplementary Figure 10. Normalization schemes for 16S data and effects on whole community data.** a–c) Strain relative abundance from 16S sequencing at $t_0$ in Exp B plotted against three technical quantities: 16S rRNA gene copy number (a), strain-specific OD-to-CFU conversion factor (b), and the product of the two (c). Each point is one strain, colored as in the legend in (e). Grey lines are ordinary least-squares fits, coefficients of determination and two-sided *p*-values are inset. No factor is significantly associated with initial relative abundance ($R^2 \leq 0.02$, $p > 0.6$ in all cases), indicating that neither copy number nor culture-density conversion accounts for the observed 16S sequencing skew. d) Measured strain composition at the final passage (passage 5) for Exp A and Exp B under possible normalizations: uncorrected amplicon read fractions ("raw"), 16S copy-number correction, and correction by a per-strain $t_0$ deviation factor (see Supplementary Methods 2, Equation [S16] for details). Both experiments converge on a low-diversity community dominated by *Arthrobacter* (Exp A) or *Burkholderia* (Exp B) together with *Mucilaginibacter* and *Rhizobium*, and this pattern is preserved across normalizations. e) Simulated strain composition at the final passage under five initialization schemes: equal initial abundance, equal initial abundance with leakage set to 0 (no cross-feeding), measured $t_0$ 16S abundances, equal OD with strain-specific OD-CFU conversion, and measured $t_0$ 16S abundances with OD-CFU conversion. Colors as in the legend. The simulation without cross-feeding is by far the most divergent outcome here and the other normalizations have only a marginal affect on the final composition.

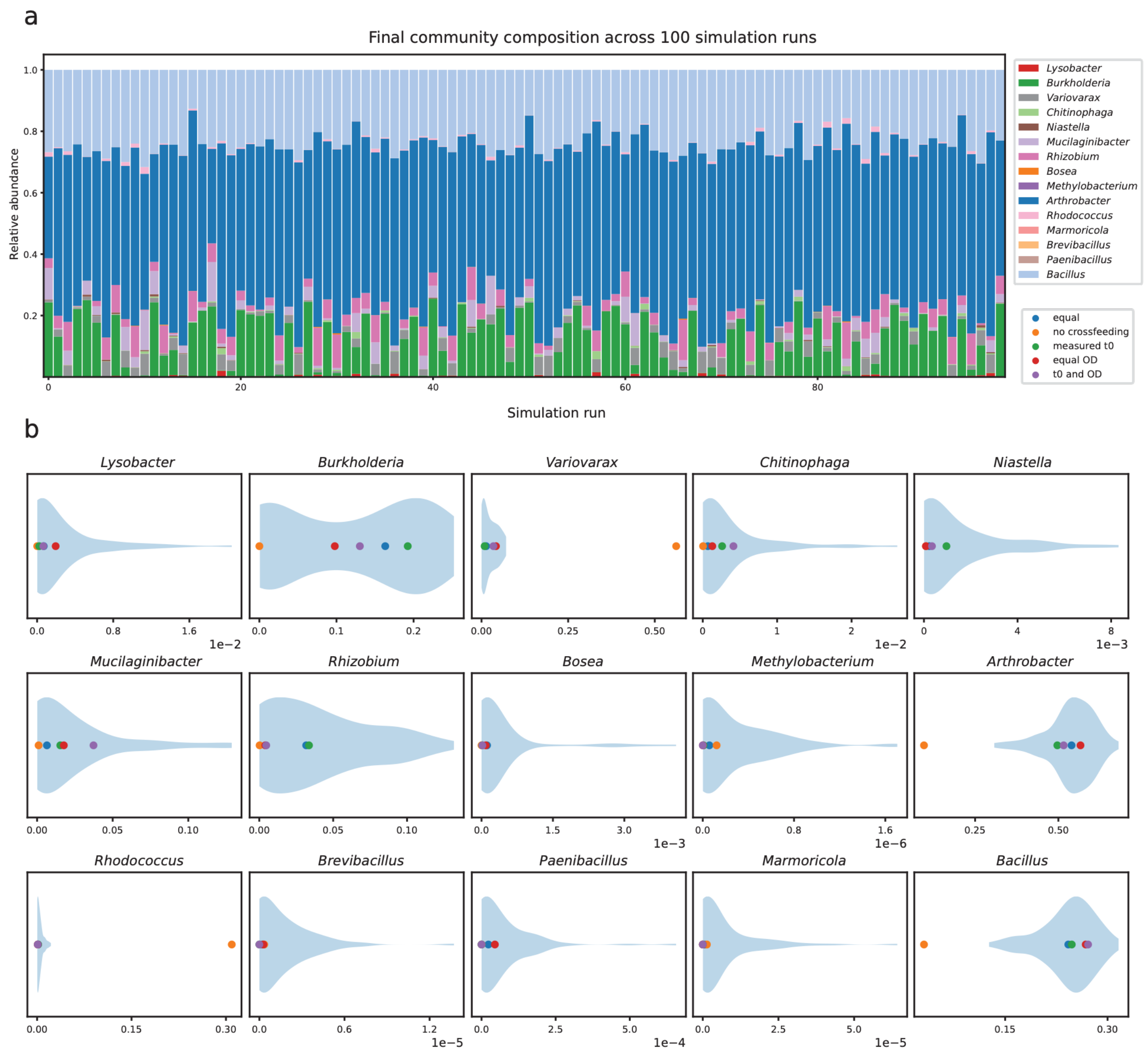


**Supplementary Figure 11. Sensitivity analysis of simulated whole community experiment with varied initial conditions.** a) Final community compositions for 100 simulation runs with varying initial community compositions. b) Distributions of strain final relative abundances from simulations in light blue with final abudances for normalization factors and simulation conditions plotted.

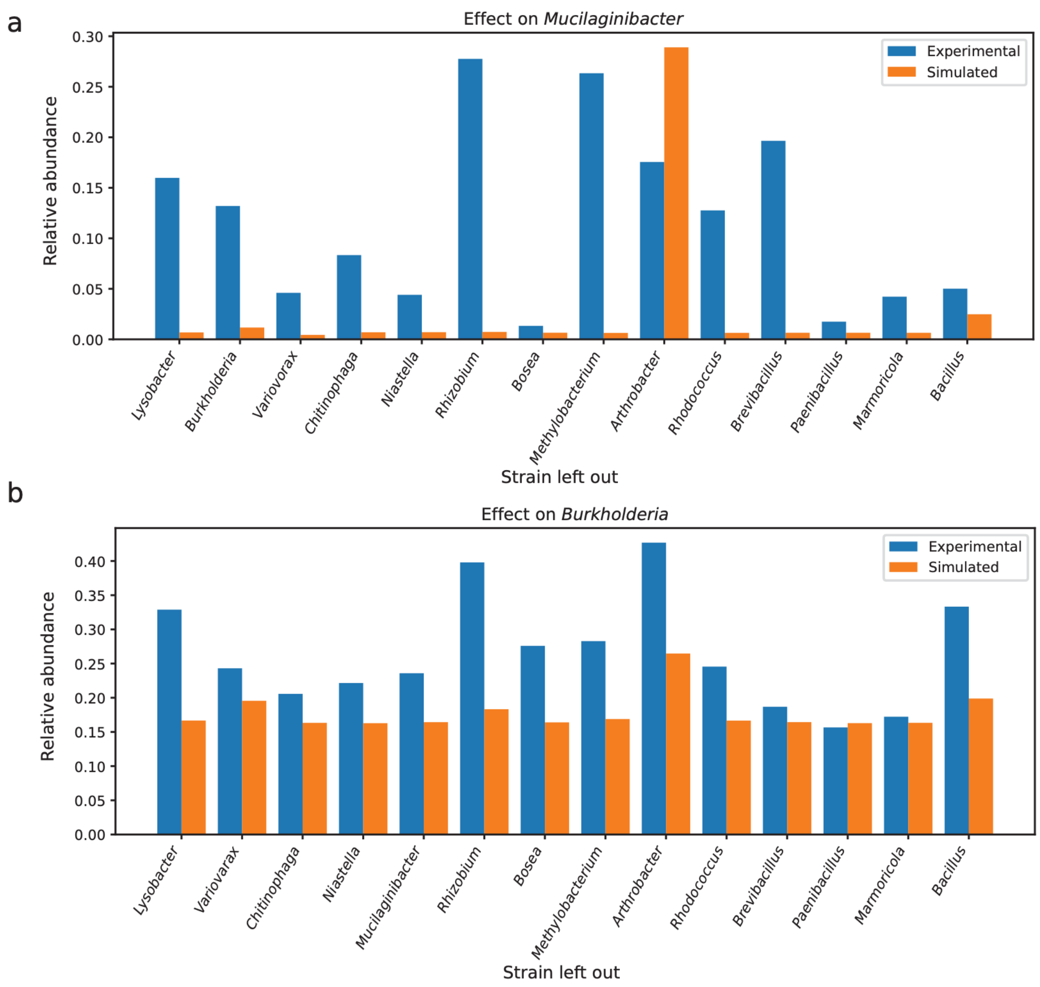


**Supplementary Figure 12. Single leave-out experiments reveal pairwise sensitivities between strains.** a) Comparison of *Burkholderia* relative abundance in experiments (blue) and CRM simulations (orange) where one strain was left out from the community. b) Comparison of *Mucilanginibacter* relative abundance in experiments (blue) and CRM simulations (orange) where one strain was left out from the community.

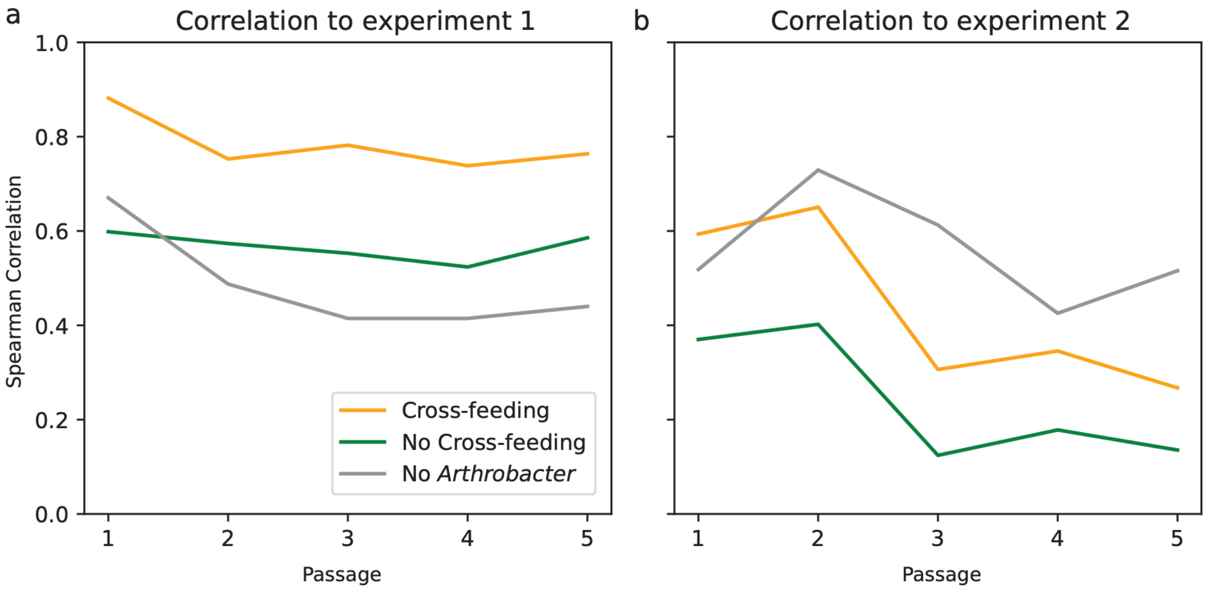


**Supplementary Figure 13. Correlations of whole community experiments to simulated communities, with *Arthrobacter* left out.** The correlation (Spearman's coefficient) between the abundance ratio of the two whole-community experiments and *Arthrobacter* leave-out (gray) or whole-community CRM simulations with cross-feeding (yellow) or without cross-feeding (green). Cross-feeding was enabled for the *Arthrobacter* leave-out simulation.

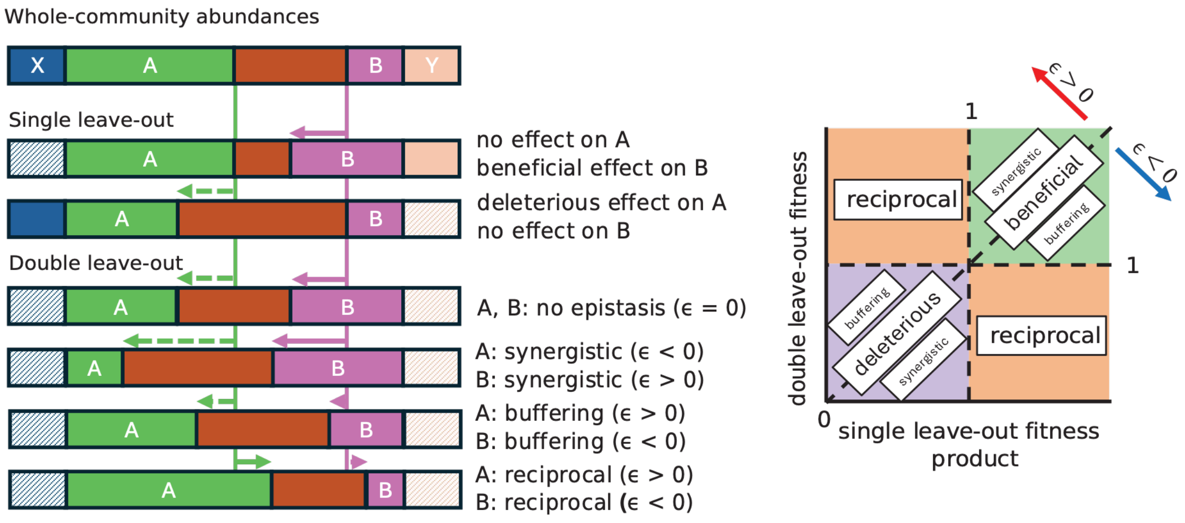


**Supplementary Figure 14. Schematic demonstrating categorization of epistatic interactions.** Here, we examine the leave-out effects of strain X,Y and how they impact relative abundance of A and B, normalized to the whole-community relative abundances. Four double leave-out scenarios are presented, resulting in different epistatic interactions. We can then examine epistatic interactions on axes relating the product of single leave-out fitness to the double leave-out fitness, especially in relation to whole-community fitness (normalized to 1).

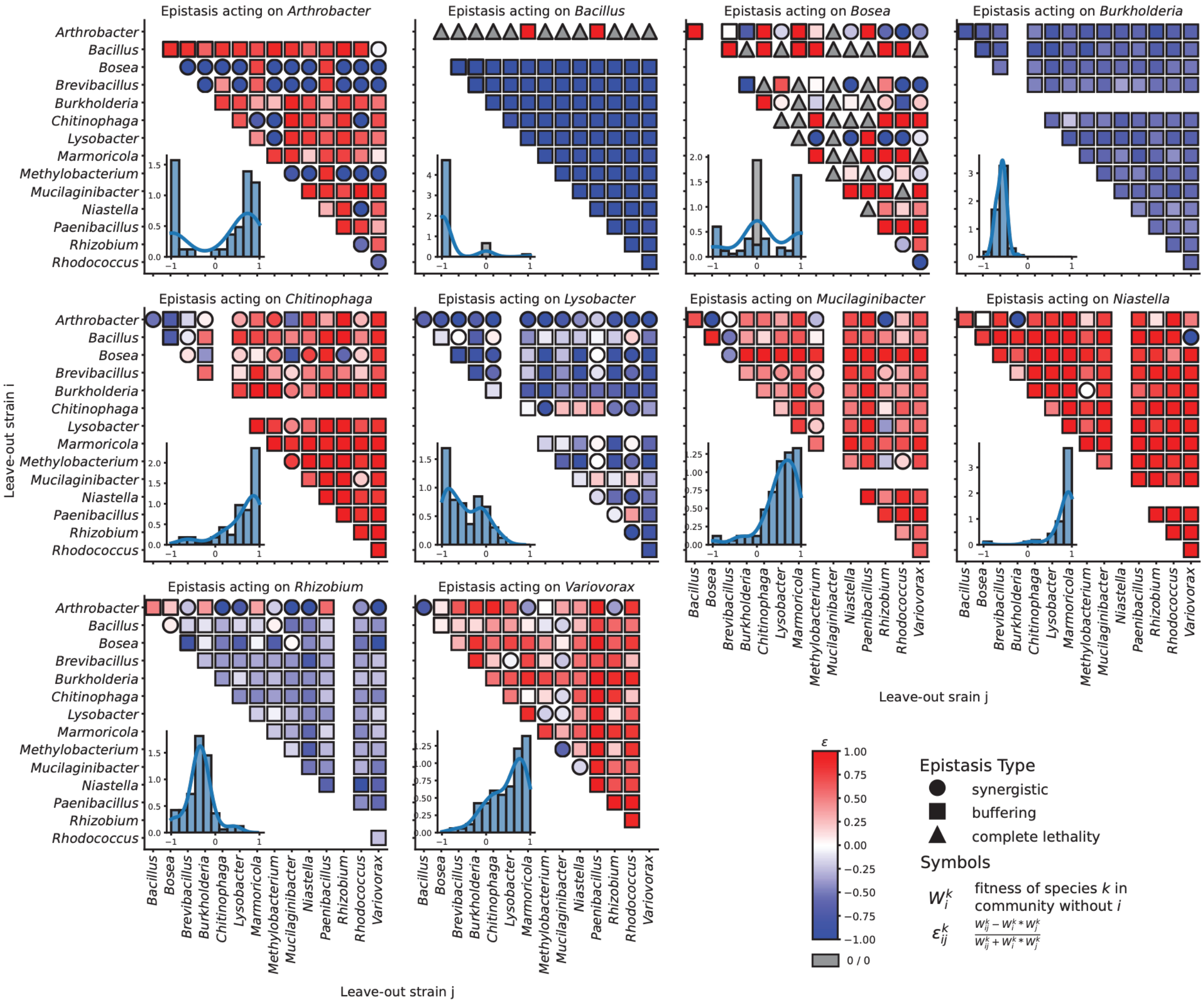


**Supplementary Figure 15. Pairwise epistatic interaction effects on relative abundance of each strain.** Each pair is colored by the epistasis value, where blue pairs exhibit decreased fitness from linear expectation, and red pairs exhibit increased fitness from linear expectations. Circles indicate synergistic nonlinear effect on relative abundance, while squares signify a buffering nonlinear impact on relative abundance. Triangles indicate that the double leave-out and at least one single leave-out resulted in zero growth for the given strain.
